# Imaging tools generated by CRISPR/Cas9 tagging reveal cytokinetic diversity in mammalian cells

**DOI:** 10.1101/2022.03.14.484313

**Authors:** Mathieu C. Husser, Imge Ozugergin, Tiziana Resta, Vincent J. J. Martin, Alisa J. Piekny

**Affiliations:** Biology Department, Concordia University, Montreal, Quebec, Canada; Center for Applied Synthetic Biology, Concordia University, Montreal, Quebec, Canada; Center for Microscopy and Cellular Imaging, Concordia University, Montreal, Quebec, Canada

## Abstract

Cytokinesis is required to physically separate the daughter cells at the end of mitosis. This process occurs via the ingression of an actomyosin ring that assembles in anaphase and pulls in the overlying plasma membrane as it constricts. Mechanistic studies have uncovered different pathways that regulate the assembly and position of the ring in mammalian cells, but the majority of these studies were done using HeLa cells with overexpressed transgenes, and the relative requirement for these mechanisms among the majority of cell types is not known. Here, we used CRISPR/Cas9 gene editing to endogenously tag cytokinesis proteins, anillin, Ect2 and RhoA, as well as other cellular components, with fluorescent proteins. These tools enabled the visualization of cytokinesis by live imaging to quantitatively study these proteins at endogenous levels. As a proof-of-concept, tagging anillin in multiple mammalian cell lines revealed cytokinetic diversity, which will be useful for studies of how mechanisms controlling cytokinesis vary among cell types. We also successfully tagged multiple cellular components in the same cell line, demonstrating the versatility of these tagging tools.

## Introduction

Cytokinesis is the physical separation of a cell into two daughter cells that occurs at the end of mitosis. This process must be tightly spatiotemporally controlled as failure can cause changes in cell fate and developmental defects or diseases (D’Avino et al., 2015; Lacroix & Maddox, 2012). Cytokinesis occurs via the assembly and ingression of a RhoA-dependent contractile ring that constricts to pull in the overlying plasma membrane. Multiple pathways have been shown to regulate ring assembly in cultured cells and model organisms (Green et al., 2012; Husser et al., 2021; Ozugergin & Piekny, 2021; Pollard & O’Shaughnessy, 2019). These pathways ensure that active RhoA is enriched at the equatorial plane to assemble the ring. Ect2 is the guanine nucleotide exchange factor (GEF) that activates RhoA during cytokinesis, and requires binding to phospholipids and the central spindle protein Cyk4 (MgcRacGAP) for its activity (Fig. 1A; Gomez-Cavazos et al., 2020; Wolfe et al., 2009; Yuce et al., 2005). The depletion of Cyk4 or Ect2 in HeLa cells prevents the accumulation of active RhoA at the equatorial cortex and leads to cytokinesis failure (Kamijo et al., 2006; Kim et al., 2005; Nishimura & Yonemura, 2006; Yuce et al., 2005; Zhao & Fang, 2005). Active RhoA (RhoA-GTP) recruits and activates effectors, including formin and RhoA kinase (ROCK), to generate actomyosin filaments and assemble the ring (Fig. 1A; Piekny et al., 2005; Pollard & O’Shaughnessy, 2019). Anillin is also recruited by active RhoA and acts as a scaffold protein that tethers the ring to the plasma membrane (Green et al., 2012; Piekny & Glotzer, 2008; Piekny & Maddox, 2010; Sun et al., 2015). In support of its crosslinking function, the depletion of anillin in HeLa or S2 cells leads to oscillations of the ring and cytokinesis failure (Hickson & O’Farrell, 2008; Piekny & Glotzer, 2008; Piekny & Maddox, 2010). Anillin may also be involved in the retention of active RhoA at the equatorial cortex, as well as its removal during constriction (Budnar et al., 2019; Carim et al., 2020; El Amine et al., 2013). Multiple mechanisms have been shown to control this core cytokinesis machinery in different model systems, but it is not known how their requirement varies among diverse cell types in any metazoan (Husser et al., 2021).

**Figure 1.**
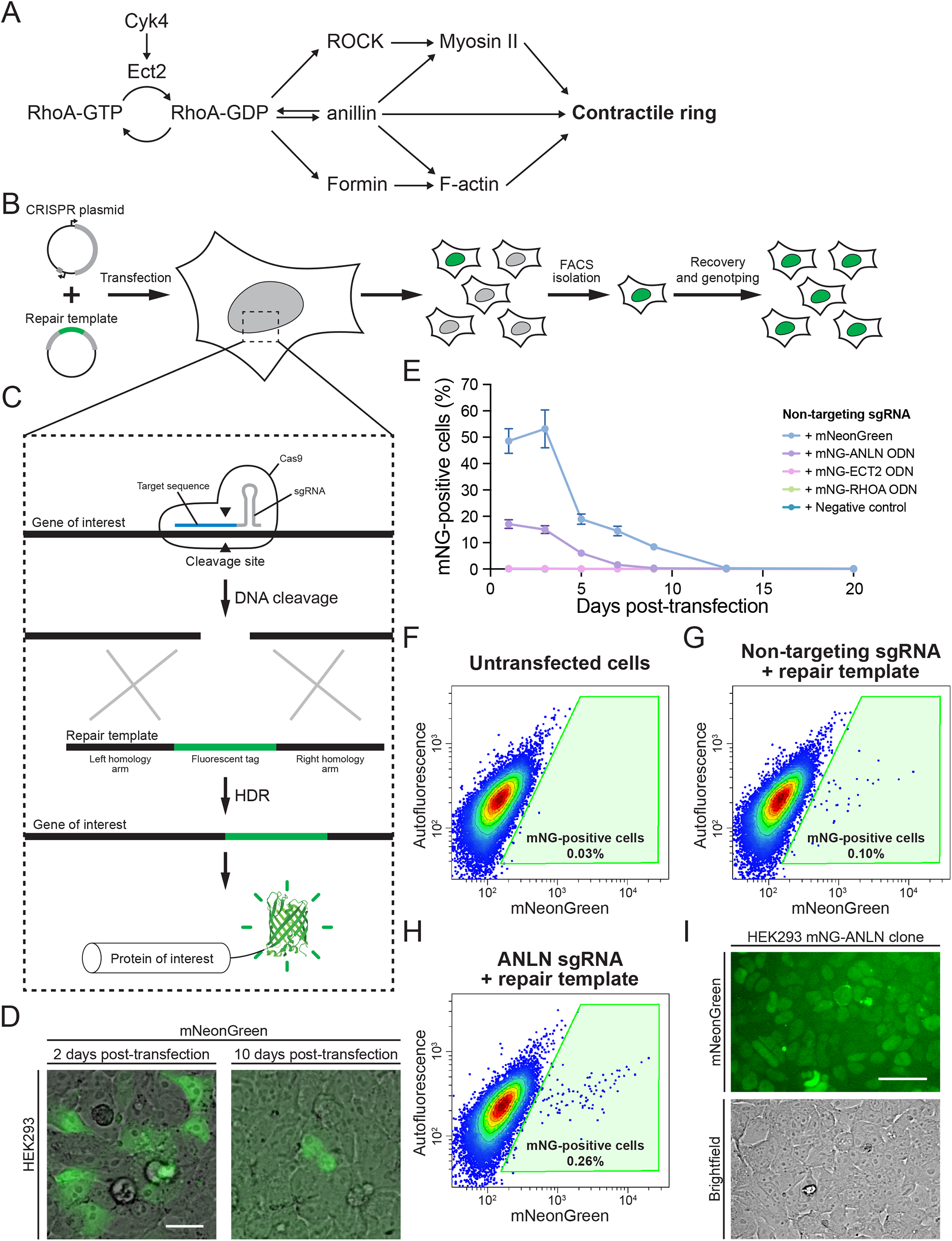
Endogenous tagging of cytokinesis using CRISPR/Cas9. A) A diagram shows the core pathway regulating contractile ring assembly during cytokinesis. The generation of active RhoA (RhoA-GTP) by its GEF Ect2 at the equatorial cortex is required for ring assembly. Active RhoA recruits and/or activates effectors (anillin, formin and ROCK proteins) to assemble actomyosin filaments into the contractile ring. B) Experimental workflow used for endogenous tagging. Cultured cells were transfected with a CRISPR plasmid and a repair template to express the CRISPR/Cas9 components and integrate a fluorescent marker. Some cells started to express the fluorescent marker after several days and were isolated by FACS. Clonal cell lines were then recovered and genotyped to verify the edits. C) Mechanism of endogenous tagging by CRISPR/Cas9. A gene-specific sgRNA directs DNA cleavage by Cas9 at the target site. The DSB generated by Cas9 can be repaired by HDR using the provided repair template and integrate a fluorescent marker at the target site. This will result in the expression of the fluorescent marker fused to the target protein. D) Representative images of ectopic mNeonGreen expression 2 days after transfection (left) and nuclear mNeonGreen-anillin signal 10 days after transfection (right). Scale bar is 25 μm. E) Percentage of mNeonGreen-positive cells over time after transfection with different mNeonGreen repair templates assessed by flow cytometry. A constitutive mNeonGreen expression vector was used as a positive control compared with the mNeonGreen-anillin, mNeonGreen-Ect2 and mNeonGreen-RhoA repair templates, and an empty plasmid as a negative control. F-H) Representative flow cytometry plots show mNeonGreen fluorescence in HEK293 cells 13 days after transfection with constructs designed to tag anillin with mNeonGreen. Non-transfected cells (F) were used to gate out non-fluorescent cells. A negative control lacking a sgRNA (G) showed residual ectopic expression of mNeonGreen from the transfection of a repair template alone. Cells transfected with both the ANLN-targeting CRISPR plasmid and repair template (H) showed a higher proportion of mNeonGreen-positive cells. The gate shown in green was used to isolate tagged cells by FACS. I) Representative image of a colony of mNeonGreen-anillin tagged HEK293 cells after single-cell isolation and recovery. Scale bar is 50 μm.

Both spindle-dependent and -independent pathways regulate cytokinesis. The central spindle recruits and activates Ect2 in proximity to the equatorial cortex where it generates active RhoA (Adriaans et al., 2019; Basant et al., 2015; Burkard et al., 2009; Frenette et al., 2012; Kotynkova et al., 2016; Lekomtsev et al., 2012; Mishima et al., 2002; Mishima et al., 2004; Petronczki et al., 2007; Su et al., 2011; Wolfe et al., 2009; Yuce et al., 2005). Astral microtubules also define the cleavage plane by directly or indirectly removing cortical regulators from the poles (Chen et al., 2021; Lewellyn et al., 2010; Mangal et al., 2018; Tse et al., 2011; van Oostende Triplet et al., 2014; Zanin et al., 2013). Spindle-independent pathways polarize the cortex via signals associated with chromatin, centrosomes and kinetochores (Beaudet et al., 2017; Beaudet et al., 2020; Cabernard et al., 2010; Canman et al., 2003; Canman et al., 2000; Kiyomitsu & Cheeseman, 2013; Ozugergin et al., 2022; Ozugergin & Piekny, 2021; Rodrigues et al., 2015; von Dassow et al., 2009; Zanin et al., 2013). A kinetochore-derived pathway induces relaxation of the polar cortices in HeLa and Drosophila S2 cells via the dephosphorylation of ERM proteins by a kinetochore-tethered protein phosphatase 1 (PP1; Rodrigues et al., 2015). Several studies also revealed a role for chromatin-associated active Ran (Ran-GTP; Ras-related nuclear protein) in coordinating the position of cortical regulators with segregating chromosomes (Beaudet et al., 2017; Beaudet et al., 2020; Kiyomitsu & Cheeseman, 2013; Ozugergin et al., 2022; Ozugergin & Piekny, 2021). All of these pathways also act in concert with the negative regulator of RhoA, MP-GAP (M-phase GTPase-activating protein), which is globally localized (Zanin et al., 2013). Having multiple mechanisms to control the function of cortical regulators ensures robust cytokinesis, but it is not clear how their requirement differs with cell type.

The mechanisms regulating cytokinesis vary widely depending on the organism and tissue (Cabernard et al., 2010; Davies et al., 2018; Fotopoulos et al., 2013; Lewellyn et al., 2010; Mangal et al., 2018; Ozugergin et al., 2022; Rodrigues et al., 2015; van Oostende Triplet et al., 2014). For example, anillin is essential for cytokinesis in cultured cells (HeLa and Drosophila S2 cells), but not in the early *Caenorhabditis elegans* zygote, and Dalmatian dogs carrying an anillin truncation mutant did not have obvious cell division defects (Hickson & O’Farrell, 2008; Holopainen et al., 2017; Maddox et al., 2005; Piekny & Glotzer, 2008). However, later in *C. elegans* development, neuronal precursor cells require anillin for cytokinesis (Fotopoulos et al., 2013; Wernike et al., 2016). Thus, the mechanisms controlling cytokinesis likely vary with parameters including fate, ploidy and geometry. In the two-cell *C. elegans* embryo, the somatic AB and germline P1 cells have different ring assembly and ingression kinetics with different levels of myosin in the ring (Ozugergin et al., 2022). In the four-cell *C. elegans* embryo, the ABa and ABp cells have stronger requirements for formin-derived F-actin compared to EMS or P2 cells, which are regulated by cell-extrinsic and intrinsic factors, respectively (Davies et al., 2018). These studies highlight the need to investigate cytokinetic diversity and understand how the mechanisms regulating cytokinesis vary among cell types.

Gene editing tools provide an opportunity to study proteins in diverse cell types (Husser et al., 2021). In human cells, cytokinesis has mostly been studied using overexpressed transgenes fused to fluorescent tags for visualization and/or affinity tags for biochemical assays. In HeLa cells, the localization of endogenous anillin fixed and stained with antibodies is similar to anillin over-expression. However, both Ect2 and RhoA show more variability and/or cause cytokinesis phenotypes when over-expressed (Chalamalasetty et al., 2006; Piekny & Glotzer, 2008; Yuce et al., 2005). TCA-fixation-based immunofluorescence microscopy is still used as one of the more reliable methods to visualize the enrichment of RhoA at the equatorial cortex (Koh et al., 2021; Schneid et al., 2021; Yonemura et al., 2004; Yuce et al., 2005), while Ect2 overexpression can lead to cytokinesis failure (Chalamalasetty et al., 2006). In addition, measurements can be confounded by the variability in expression between transfected cells, and endogenous probes enable more quantitative measurements (Husser et al., 2021). Studies have shown that proteins tagged endogenously with fluorescent proteins report more accurately and cause fewer phenotypes than using overexpressed transgenes (Mahen et al., 2014). Moreover, the same tools can be used to introduce genetic edits into different cell lines derived from the same organism.

The most popular tool for gene editing is CRISPR/Cas9, which is comprised of a Cas9 nuclease and a sgRNA (Single guide RNA; Fig. 1B, C; Cong et al., 2013; Mali et al., 2013; Pickar-Oliver & Gersbach, 2019; Wang et al., 2016). The sgRNA contains a 20-nucleotide target sequence that corresponds to a genomic target site. Cas9 is targeted to this site by the sgRNA and cleaves the DNA to introduce a double-stranded break (DSB). Human cells typically repair DSBs by non-homologous end joining (NHEJ), but can also use the homology-directed repair (HDR) pathway, which makes use of a homologous repair template to fill in the gap where the DSB was introduced (Scully et al., 2019; Wright et al., 2018). To introduce a fluorescent marker at a precise location in the genome of human cells, CRISPR/Cas9 can be used along with a synthetic repair template designed to carry the fluorescent marker flanked with homology arms for HDR (Fig. 1C; Verma et al., 2017). When introduced in frame with a gene, the fluorescent marker will be expressed fused to the protein of interest. Efforts to share validated tools for endogenous tagging (sgRNA sequence and repair template) have made these tags more readily accessible and easy to use (de Man et al., 2021; Iyer et al., 2019; Pinder et al., 2015; Roberts et al., 2017; Sakuma et al., 2016; Savic et al., 2015; Sun et al., 2021; Allencell.org; Addgene.org). However, despite these shared resources and the advantages to using endogenous tags, few cytokinesis proteins have been tagged endogenously in human cells (Mahen et al., 2014; Peterman et al., 2020).

In this work, we expanded on the set of available tools to introduce endogenous tags by generating constructs to tag anillin, Ect2 and RhoA, and by re-purposing existing constructs to tag cellular markers with different fluorophores. We used CRISPR/Cas9 to generate HeLa cell lines where fluorescent proteins were endogenously fused to anillin, Ect2 and RhoA and compared their spatiotemporal distribution during cytokinesis. We also tagged anillin in multiple human and mammalian cell lines and found differences in the breadth and timing of anillin’s cortical localization during cytokinesis. Finally, we explored the potential of this toolkit by generating cell lines that carry multiple endogenous tags of different colour. By making these tools available to the cell biology community, we hope to fuel new studies of the mechanisms regulating cytokinesis in diverse human cell types.

## Results

### Endogenous tagging of cytokinesis proteins

First, we generated tools to endogenously tag anillin (ANLN), Ect2 (ECT2) and RhoA (RHOA) with mNeonGreen in human cells using CRISPR-Cas9 and HDR (Fig. 1A-C). We designed sgRNAs to target these genes, and repair templates to integrate the coding sequence of mNeonGreen in frame with the coding sequence of the ANLN, ECT2 and RHOA genes. The repair templates were designed to have 1kb homology arms flanking the mNeonGreen gene. Three sgRNAs were designed for each gene (Table S1) and cloned into a plasmid expressing the high-specificity HypaCas9 protein. We only targeted the N-terminus of RhoA because its C-terminus is post-translationally cleaved, but we targeted both ends of the ANLN and ECT2 genes for tagging. The sets of sgRNA-containing CRISPR plasmids and repair templates were validated by co-transfecting them into HEK293 (immortalized human embryonic kidney cells, female) cells and monitoring the appearance of a mNeonGreen signal with expected localization patterns.

When generating endogenously tagged cell lines, we observed non-specific ectopic expression of mNeonGreen directly following transfection (Fig. 1D). This ectopic signal has been reported previously (Fueller et al., 2020), and likely arises from ectopic expression from the repair template rather than genomic integration, as it progressively disappears over ∼9 days following transfection (Fig. 1E). Thus, to avoid mistaking ectopic expression for the endogenous tag, we recommend culturing the cells for 8 days after transfection to allow for the ectopic signal to decrease and the endogenous tag to be expressed. We observed the expected localization pattern for the tagged proteins starting ∼4 days after transfection.

Using the workflow in Fig. 1B, we generated N-terminal mNeonGreen fusions with anillin, Ect2 and RhoA in HEK293 cells. Notably, we did not recover cells with C-terminal tags fused to anillin, while cells with C-terminal tags fused to Ect2 did not show the correct cellular localization (data not shown). The C-terminal tag likely disrupts the function of Ect2, but it is not clear why the editing efficiency was insufficient for tagging the C-terminal end of anillin. The successful repair templates and sgRNAs used for each loci are listed in Table 1. After confirming tagging by visual inspection, single fluorescent cells were isolated by FACS (fluorescence-activated cell sorting; e.g. Fig. 1F-H). Single-cell clones were screened by fluorescence microscopy (e.g. Fig. 1I), PCR and sequencing (e.g. Fig. S1) to validate the cell lines. Finally, this workflow was repeated to generate HeLa (human cervical carcinoma, female) cell lines expressing mNeonGreen-tagged anillin, Ect2 and RhoA. We were able to isolate HEK293 and HeLa cell lines where anillin and Ect2 were homozygously tagged. However, we were unable to isolate homozygously tagged RhoA cell lines despite screening 72 clones of RhoA-tagged HEK293 and 24 clones of RhoA-tagged HeLa. This may be due to the low tagging efficiency at this locus, or some impediment to RhoA function caused by the tag.

**Table 1.**
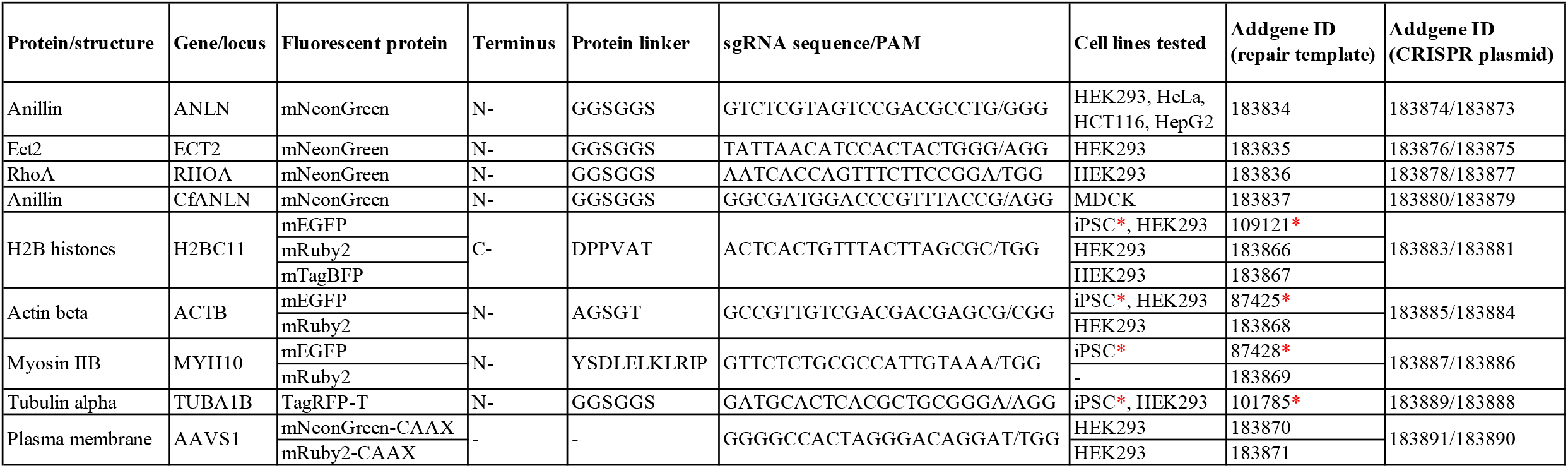
A toolkit of repair templates to tag multiple proteins and cellular components. The repair templates generated and/or used in this study are listed. For each protein or component, the terminus targeted for tagging, the protein linker sequence, and the different fluorophores that are available are indicated. The best sgRNA sequence used for CRISPR targeting of each locus is also indicated. The cell lines that each repair template has been tested in and the Addgene ID for each repair template are also listed. Red asterisks indicate repair templates that were created and tested in WTC-11 iPSCs by the Allen Institute for Cell Science (Obtained from Addgene; Roberts et al., 2017; Allencell.org).

### Visualization of endogenously tagged cytokinesis proteins

We characterized the localization of endogenous mNeonGreen-anillin, -Ect2 and -RhoA in HeLa cells by fluorescence microscopy. First, we compared the expression of the endogenous tags to transiently transfected transgenes. When comparing fields of view of the endogenously tagged cells to the overexpressed transgenes, there was a striking difference in variability and levels across the cell populations. The endogenously tagged proteins were more consistent in their expression and were expressed at much lower levels compared to the overexpressed transgenes (Fig. 2A, B). In the endogenously tagged cell lines, anillin and Ect2 were nuclear in interphase cells as expected, while RhoA was cytosolic (Fig. 2A). Also, while there was some variability in anillin and Ect2 expression, this was likely due to their cell cycle-dependent turnover and was far less than what was observed for the transgenes (Fig. 2A, B). Thus, endogenous probes should be useful to accurately measure protein expression and localization.

**Figure 2.**
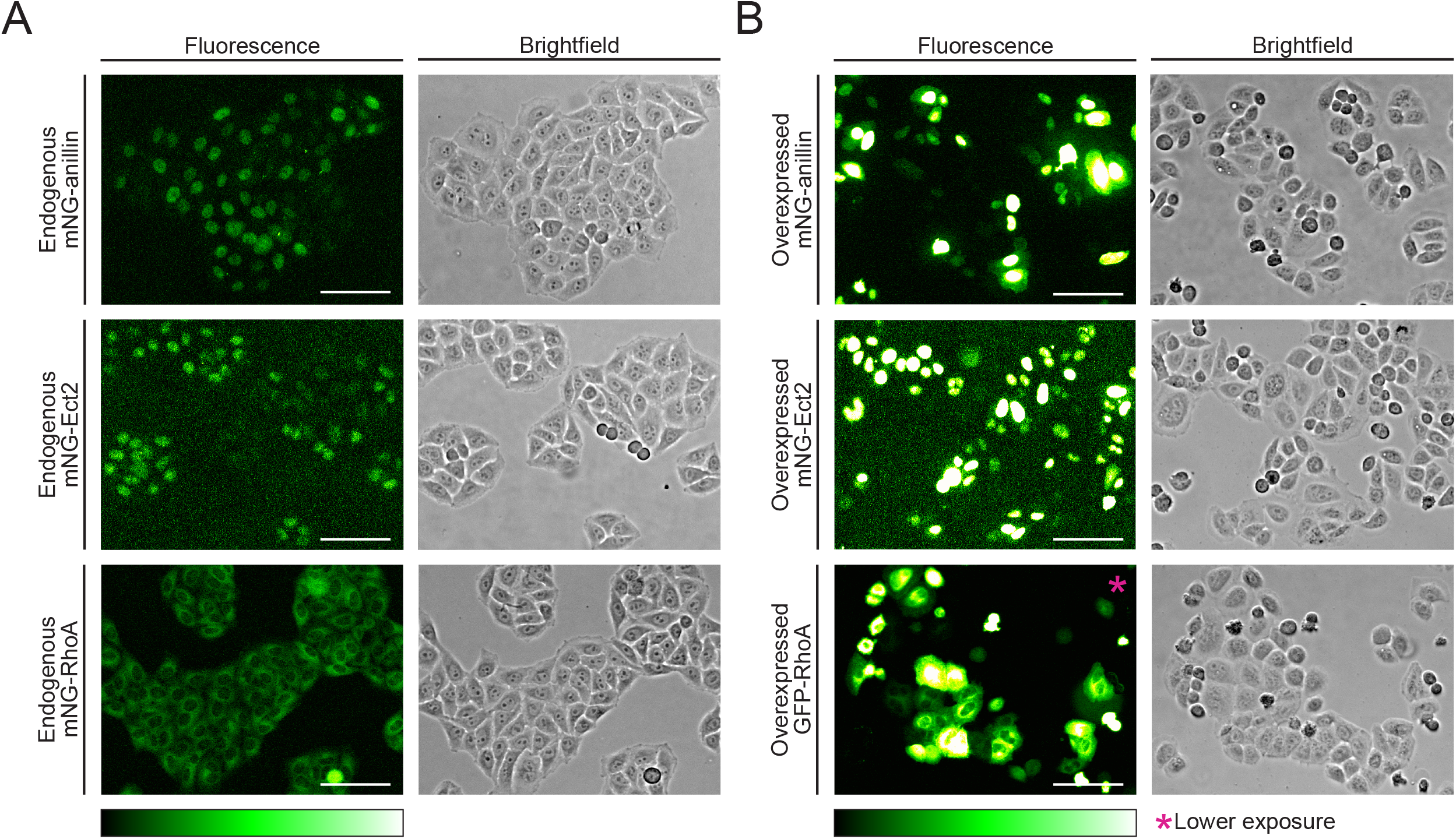
Comparison of endogenous tags with transient overexpression. A) Fluorescent (left) and corresponding brightfield (right) images of HeLa cells where anillin (top), Ect2 (middle) and RhoA (bottom) are endogenously tagged with mNeonGreen. B) Fluorescent (left) and corresponding brightfield (right) images of HeLa cells where mNeonGreen-anillin (top), mNeonGreen-Ect2 (middle) and GFP-RhoA (bottom) are exogenously expressed, 24 hours after transfection. *Image taken with a lower exposure time. Scale bars are 100 μm.

We then imaged HeLa cells during cytokinesis to follow the localization of endogenously tagged anillin, Ect2 and RhoA during cytokinesis (Fig. 3A-C). As expected, anillin was enriched at the equatorial cortex shortly after anaphase onset and remained in the furrow throughout ingression, after which it localized to the midbody (Fig. 3A). We measured this enrichment by using a linescan to plot the intensity of mNeonGreen-anillin along the cortex of cells at furrow initiation (∼9-14 min; Fig. 3D, and E, left). To visualize localization over time, we repeated this analysis every two minutes, starting two minutes before anaphase onset (Fig. 3F and G, left). We observed a gradual decrease in the breadth and an increase in the enrichment of anillin at the equatorial cortex over time, with the enrichment being first visible 4-6 minutes after anaphase onset (Fig. 3G, left). We similarly imaged Ect2 during cytokinesis (Fig. 3B). Ect2 initially localized to the central spindle, then was also visible at the equatorial cortex and remained in both locations through ingression, followed by localization to the midbody (Fig. 3B and S2A, B). We used linescans to measure the intensity of mNeonGreen-Ect2 along the cortex at furrow initiation (Fig. 3E, middle), and along the cortex and central spindle over time (Fig. 3G, middle, H and I). Ect2’s localization to the central spindle preceded the equatorial cortex, which was first visible ∼6 minutes after anaphase onset and was narrow compared to anillin. Lastly, we found that RhoA was also enriched at the equatorial cortex at the onset of ingression (Fig. 3C, E, right). Linescans revealed that this enrichment was visible ∼6-8 minutes after anaphase onset (Fig. 3C, G, right). This enrichment appeared weak compared to Ect2 or anillin, likely because of the large pool of cytoplasmic RhoA. To further demonstrate that mNeonGreen-RhoA is functional, we tested its ability to respond to GEF activity by transiently overexpressing the C-terminus of Ect2 [Ect2 (C-term); amino acids 420-882], which contains the DH domain required for nucleotide exchange and was previously shown to ectopically activate RhoA (Yuce et al., 2005). Indeed, we observed an increase in cortical localization of mNeonGreen-RhoA in interphase cells expressing mScarlet-I-Ect2 (C-term) (Fig. S2C, D; Yuce et al., 2005).

**Figure 3.**
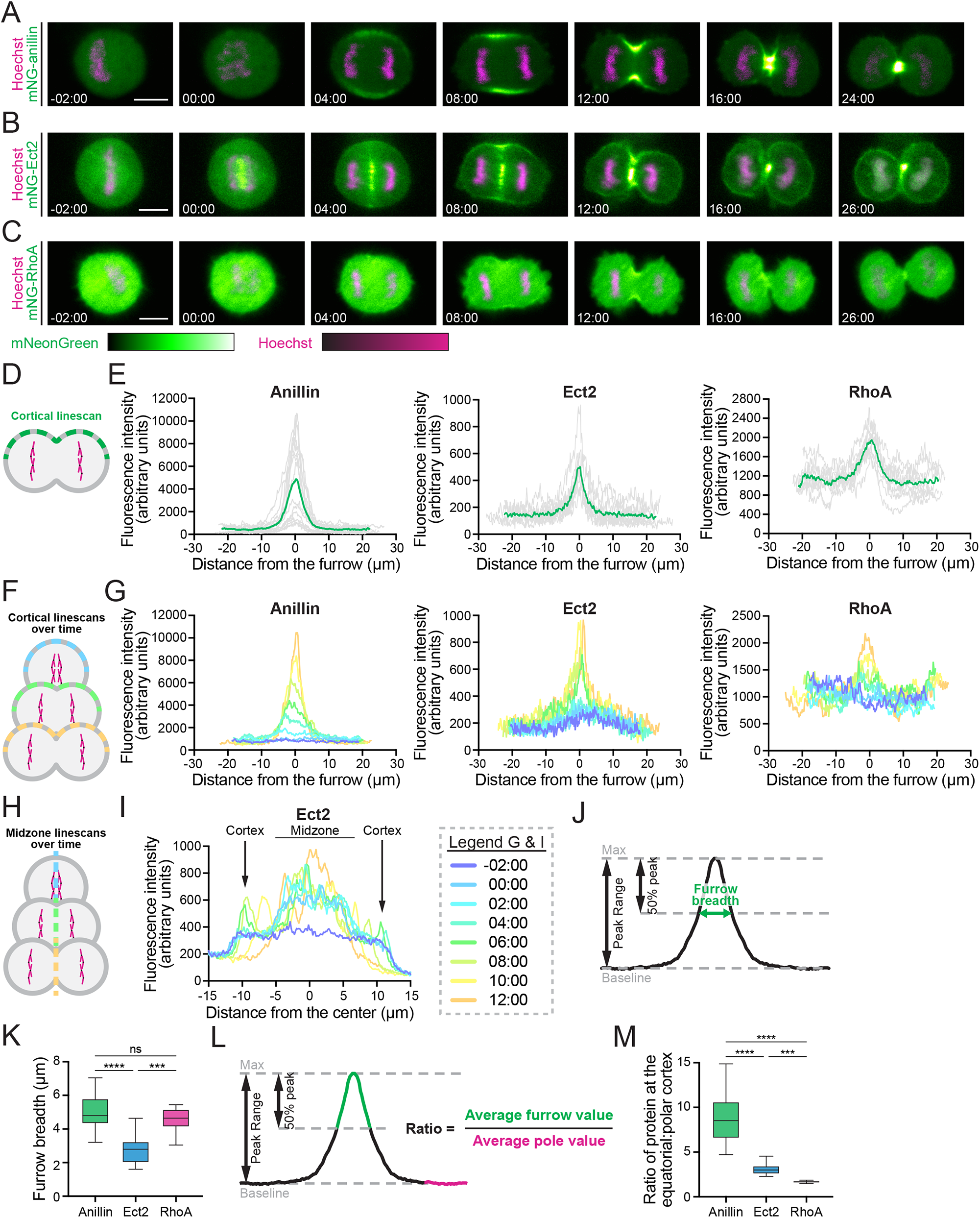
Comparison of endogenous anillin, Ect2 and RhoA localization in HeLa cells during cytokinesis. A-C) Timelapse images show cells expressing endogenous mNeonGreen-anillin (A), mNeonGreen-Ect2 (B) and mNeonGreen-RhoA (C) during cytokinesis. mNeonGreen is shown in green and DNA (stained with Hoechst) in magenta. Times are shown relative to anaphase onset. The scale bar is 10 μm. D) Schematic of the linescans used to plot the fluorescent intensity along the cell cortex during the furrow initiation phase in E. E) Graphs show fluorescence intensity along the cortex of HeLa cells expressing endogenously tagged anillin (left, n=16), Ect2 (middle, n=10) and RhoA (right, n=10). Individual replicates are shown in grey and the average is shown in green. F) Schematic of the linescans used to plot the fluorescent intensity along the cell cortex over time in G. G) Graphs show fluorescence intensity along the cortex of a single HeLa cell expressing endogenously tagged anillin (left), Ect2 (middle) and RhoA (right) at multiple time points starting at 2 minutes before anaphase onset, shown in different colors as indicated in the scale below. H) Schematic of the linescans used to plot the fluorescent intensity perpendicular to the cell equator over time in I. I) A graph shows fluorescence intensity along the perpendicular axis of a single HeLa cell expressing endogenously tagged Ect2 at multiple time points starting 2 minutes before anaphase onset, shown in the same colors as G. J) A schematic shows how the breadth at the equatorial cortex was calculated for K. K) Box plots show the breadth of anillin (n=16), Ect2 (n=10) and RhoA (n=10) in HeLa cells. L) A schematic shows how the ratio of protein at the equatorial cortex relative to the polar cortex was calculated for M. M) Box plots show the enrichment of anillin (n=16), Ect2 (n=10) and RhoA (n=10) at the equatorial cortex in HeLa cells. Box plots in K and M show the median line, quartile box edges and minimum and maximum whiskers. Statistical significance was determined by one-way ANOVA (ns, not significant; * p≤0.05; ** p≤0.01; *** p≤0.001; **** p≤0.0001).

Next, we quantified the linescan measurements to demonstrate the usefulness of endogenous tags for functional studies of cytokinesis. We measured the breadth of localization of each endogenously tagged protein, along with the ratio of enrichment at the equatorial cortex relative to the polar cortex at furrow initiation (Fig. 3J-M). To measure breadth, we used the linescans to determine the number of pixels above 50% of the peak’s intensity (Fig 3J). Interestingly, the breadth of RhoA and anillin was similar, while Ect2 was more restricted (Fig. 3K). Similar results were obtained when measuring breadth in proportion to cortex length, showing that these differences are independent of cell size (Fig. S3A). To measure the accumulation of proteins in the furrow, we calculated the ratio of the average pixel intensity in the peak to the average pixel intensity at the polar cortex (Fig. 3L). As expected from expression level and breadth, anillin was most enriched in the furrow, with 8.86 +/- 2.94 times more protein in the furrow than at the poles (n=16) compared to Ect2 (3.09 +/- 0.70 fold, n=10) and RhoA (1.67 +/- 0.13 fold, n=10; Fig. 3M).

### Endogenous tagging of anillin in different mammalian cell lines

With our endogenous tagging tools, we tested our ability to tag anillin with mNeonGreen in different human and mammalian cell lines. In addition to mNeonGreen tagging of anillin in HEK293 and HeLa cells, we tagged anillin in HCT116 (human colorectal carcinoma, male), HepG2 (human hepatocellular carcinoma, male) and MDCK (madin-darby canine kidney epithelial cells, female) cells. Since MDCK are canine cells, we created new sgRNA and repair templates suited for the genome of this organism (Table 1). These cell lines presented different challenges for gene editing. Some cell types, such as HCT116 and MDCK cells had low transfection efficiency using liposome-based methods (*e*.*g*. Lipofectamine 3000), which required using nucleofection to increase the efficiency of transfection and frequency of tagged cells. We also used NHEJ inhibitors to increase the efficiency of integration by HDR. Another limitation was the rate of single-cell recovery after FACS. Some cell lines like HeLa and MDCK recovered well from single-cell isolation. However, other cell lines, like HepG2, were very recalcitrant to growing from a single cell, and we initially only recovered two non-fluorescent colonies from 120 single FACS-sorted HepG2 cells. To resolve this issue, we enriched tagged cells by sorting 15,000 fluorescent cells together as a population and allowing them to recover for a few days. Then, we isolated clones by sorting 10-25 fluorescent cells per well of a 96-well plate and monitoring for the recovery of a single colony in each well. By incorporating these troubleshooting methods into our protocols, we successfully endogenously tagged anillin with mNeonGreen in HeLa, HEK293, HCT116, HepG2 and MDCK cells.

### Cytokinetic diversity across mammalian cell lines

After tagging anillin endogenously in diverse cell types, we compared mNeonGreen-anillin localization during cytokinesis (Fig. 3A, 4A-D). First, we observed that mNeonGreen-anillin is globally localized at the cortex of HEK293, HCT116 and MDCK cells during metaphase, but not in HeLa and HepG2 cells, where it is strictly cytosolic (Fig. 3A, 4A-D). In all cell lines, mNeonGreen-anillin was first visible at the equatorial cortex ∼4-8 minutes after anaphase onset and remained in the furrow throughout ingression, after which it localized to the midbody (Fig. 3A, 4A-D). Linescans were used to measure mNeonGreen-anillin along the cortex in the different cell lines at furrow initiation (Fig. 3D, left and 4E-H), and every 2 minutes starting 2 minutes before anaphase and until mid-late closure (Fig. 3E, left and 4I-L). We also measured the duration of furrow ingression from anaphase onset to the end of ring closure in these cell lines (Fig. 4M, N). We found that the duration of ingression was similar in HeLa (18.1 +/- 1.8 min, n=14), HEK293 (19.7 +/- 4.5 min, n=11) and MDCK cells (16.0 +/- 4.3 min, n=8; Fig. 4N). HepG2 cells took the longest to ingress (26.6 +/- 5.4 min, n=13), while HCT116 cells took the least amount of time (10.8 +/- 1.1 min, n=9). The breadth and intensity of mNeonGreen-anillin in the furrow also varied among cell lines (Fig. 3D, left and 4E-H), with a broader peak in HCT116 cells and a narrower peak in HepG2 cells (Fig. 4F, G). We also observed differences in the localization of mNeonGreen-anillin over time in the different cell lines. In HCT116, HEK293 and MDCK cells where mNeonGreen-anillin was at the cortex during metaphase, these pools were removed or shifted to the equatorial cortex by 4-8 minutes after anaphase onset (Fig. 4I, J, L). Also, while the breadth of mNeonGreen-anillin in HepG2 cells remained consistently narrow, the mNeonGreen-anillin peaks were broad and became narrow with time in the other cell lines (Fig. 3E, left and 4I-L). Indeed, measurements of breadth and equatorial to polar cortical enrichment supported our observations (Fig 4O, P). mNeonGreen-anillin was broader in HCT116 cells, narrower in HepG2 cells, and more similar in HeLa, HEK293 and MDCK cells (Fig. 4O). Interestingly, the breadth appears to be inversely proportional to the duration of ingression, with HCT116 cells ingressing the fastest and HepG2 ingressing the slowest (Fig. 4N). We were surprised to see that mNeonGreen-anillin was similarly enriched in the equatorial cortex of HeLa, HEK293 and HCT116 cells (7.1 +/- 2.4, 6.3 +/- 2.3 and 6.3 +/- 1.9 fold enrichment, respectively), while HepG2 cells had a significantly stronger enrichment (10.4 +/- 2.6 fold), and MDCK cells had a weaker enrichment (3.7 +/- 1.5 fold; Fig. 4P). Similar results were obtained when measuring breadth as a percentage of cortex length, showing that these differences are independent of cell size (Fig. S3B). These data suggest that the breadth rather than accumulated levels may influence the rate of ingression. This is further supported by our observation that within our homozygous mNeonGreen-anillin HeLa cell line, there were two populations of cells with higher vs. lower levels of expression (Fig. 3E, left). This difference is likely due to heterogeneity at other loci since these cells are aneuploid. We plotted these two populations of anillin-tagged HeLa cells side-by-side and compared the same parameters (Fig. S3C-J). We found no difference in the duration of ingression and in the breadth of the furrow between the two populations (Fig. S3D-H). In addition, we observed variability in how MDCK cells ingress. MDCK cells can polarize in culture when grown to confluency and are often used to study events required for apicobasal polarity (Riga et al., 2020; Rodriguez-Boulan & Macara, 2014). Interestingly, we found that these cells displayed a variety of ingression symmetries, ranging from symmetric to highly asymmetric (Fig. S4). Since these cells were imaged when they were not confluent, these variabilities could reflect intrinsic differences, and/or extrinsic influences from neighbouring cells before polarization (Herszterg et al., 2014).

**Figure 4.**
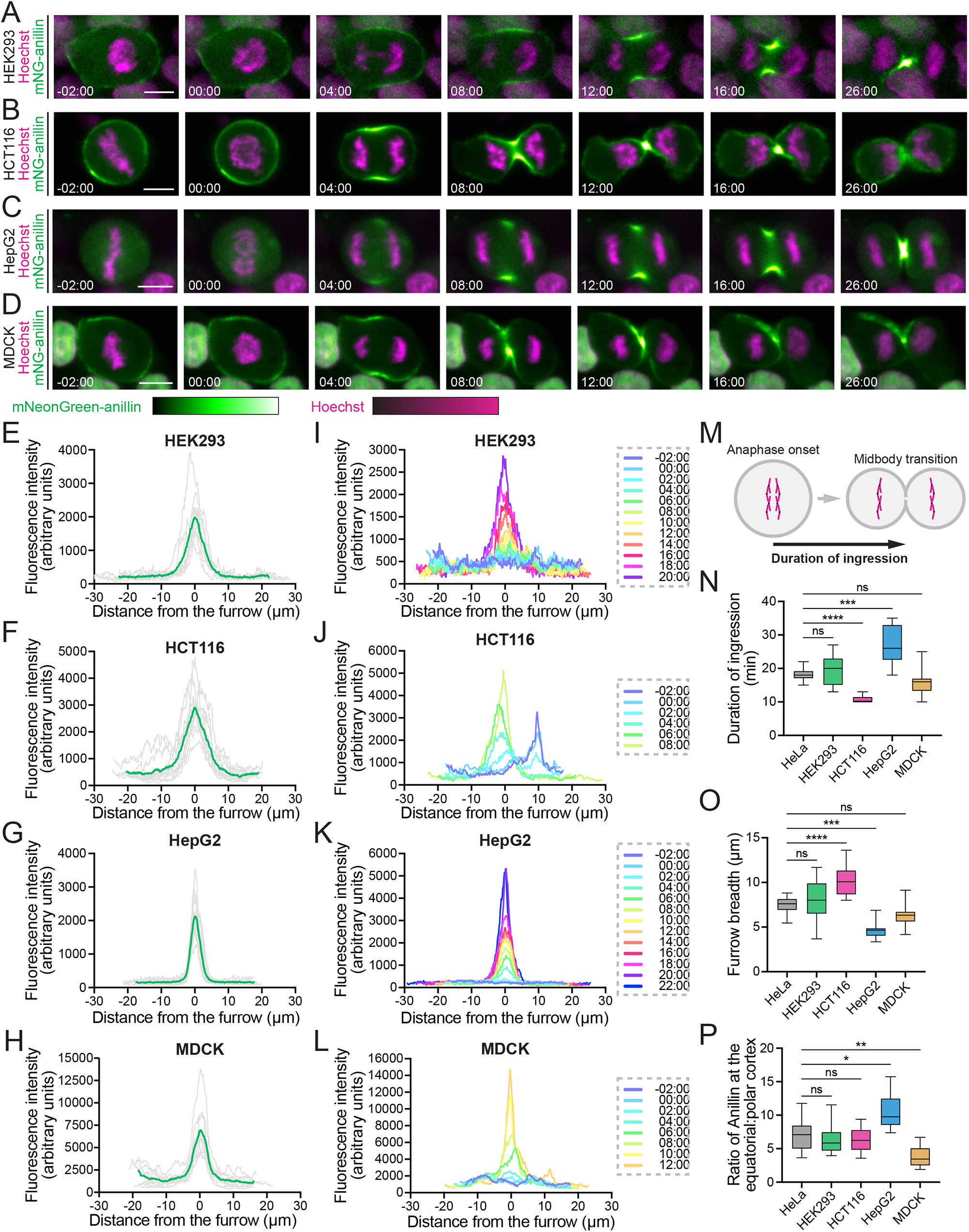
Endogenous tagging of anillin reveals cytokinetic differences between different cell lines. A-D) Timelapse images show endogenous mNeonGreen-anillin in HEK293 (A), HCT116 (B), HepG2 (C), and MDCK (D) cells during cytokinesis. mNeonGreen is shown in green and DNA (stained with Hoechst) is in magenta. Times are shown relative to anaphase onset. The scale bar is 10 μm. E-H) Graphs of linescans show fluorescence intensity of mNeonGreen-anillin along the cortex of HEK293 (E, n=9), HCT116 (F, n=11), HepG2 (G, n=13), and MDCK (H, n=10) cells at furrow initiation. I-L) Graphs of linescans show fluorescence intensity of mNeonGreen-anillin along the cortex of a single HEK293 (I), HCT116 (J), HepG2 (K) and MDCK (L) cell at multiple time points starting at 2 minutes before anaphase onset, shown in different colors as indicated by the scale. M) A schematic shows how ingression time was measured. N) A box plot shows the duration of ingression in HeLa (n=14), HEK293 (n=11), HCT116 (n=9), HepG2 (n=13) and MDCK (n=8) cells. O) A blox plot shows the breadth of mNeonGreen-anillin localization at the equatorial cortex at furrow initiation in HeLa (n=16), HEK293 (n=9), HCT116 (n=11), HepG2 (n=13) and MDCK (n=10) cells. P) A box plot shows the ratio of anillin at the equatorial cortex compared to the polar cortex in HeLa (n=16), HEK293 (n=9), HCT116 (n=11), HepG2 (n=13) and MDCK (n=10) cells. Box plots in N-P show the median line, quartile box edges and minimum and maximum whiskers. Statistical significance was determined by one-way ANOVA (ns, not significant; * p≤0.05; ** p≤0.01; *** p≤0.001; **** p≤0.0001).

### Endogenous tagging of cellular markers

Several constructs have been made available for endogenous tagging and presumably can be used in any human cell type (de Man et al., 2021; Iyer et al., 2019; Pinder et al., 2015; Roberts et al., 2017; Sakuma et al., 2016; Savic et al., 2015; Sun et al., 2021; Allencell.org; Addgene.org). However, these tools have only been validated in a few cell lines, and the choice of fluorophore is limited. To expand on this toolset, we obtained repair templates and sgRNA sequences from the Allen Institute for Cell Science (through Addgene) and replaced the mEGFP tag with either a red (mRuby2) or a blue (TagBFP) fluorophore. We also generated repair templates targeted to the AAVS1 locus to express membrane-specific tags by fusing mNeonGreen or mRuby2 with the CAAX domain of K-Ras. All of the repair templates and sgRNAs used and generated in this study are listed in Table 1. All of the constructs were validated in HEK293 cells (individual lines not shown, multi-tagged lines are shown in Figure 5). Notably, we were unable to obtain MYH10-tagged cells after several attempts using 4 different sgRNAs in HEK293 and HeLa cells. Since this protein was tagged successfully in iPSCs (Roberts et al., 2017), there could be cell-specific differences in how this gene is expressed, or the efficiency of repair.

**Figure 5.**
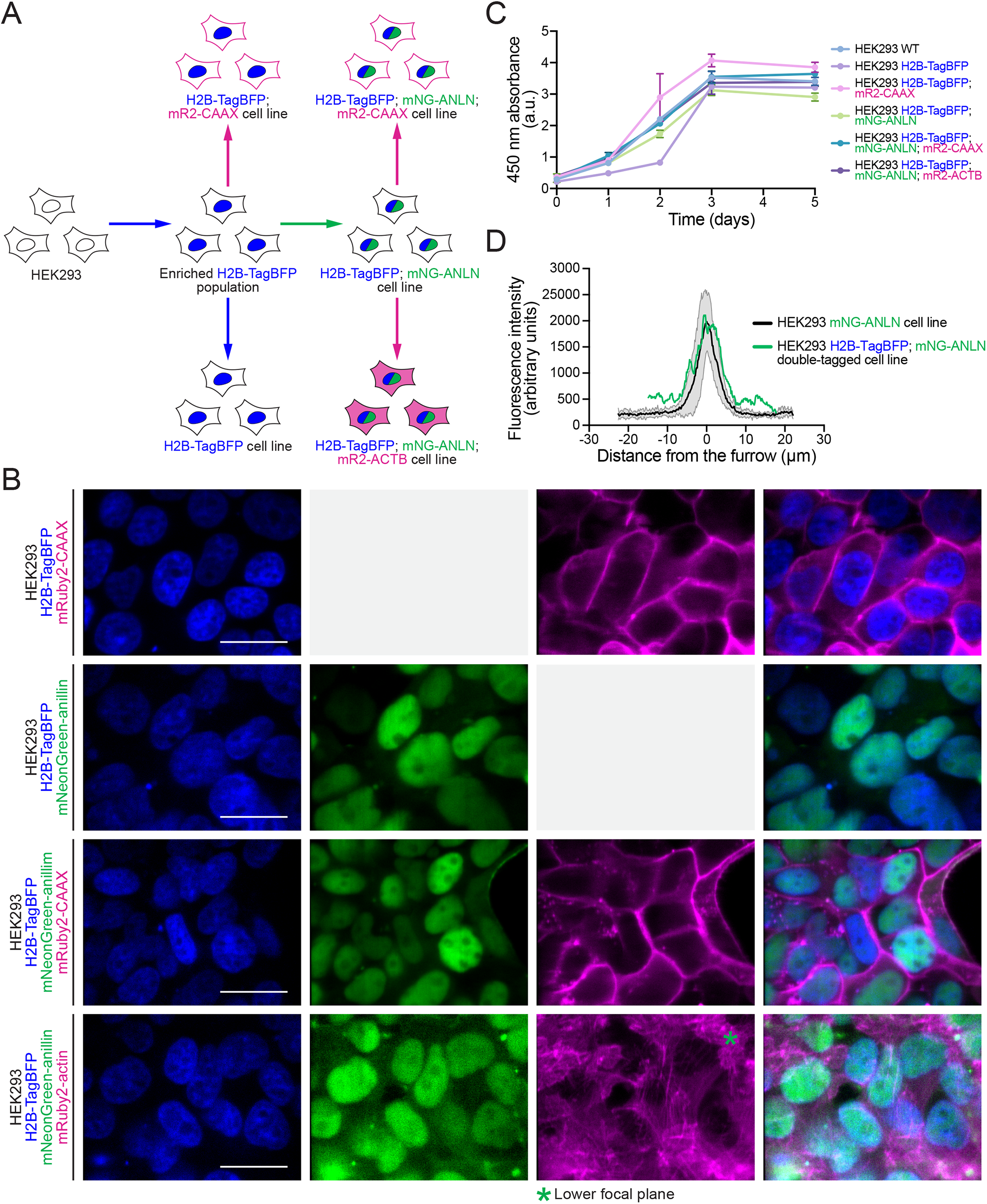
Generation of multi-tagged cell lines for multi-color fluorescence imaging. A) A schematic shows the workflow used to generate multi-tagged HEK293 cell lines. H2B histones were first tagged with TagBFP and FACS was used to obtain an ‘enriched’ population. From this enriched population, anillin or the plasma membrane were tagged with mNeonGreen and mRuby2, respectively. Clonal lines were isolated along the way to collect single and double-tagged cell lines. Finally, triple-tagged cell lines were generated and isolated by tagging β-actin or the plasma membrane with mRuby2 in the H2B-TagBFP; mNeonGreen-anillin double-tagged cell line. B) Fluorescent images show the multi-tagged HEK293 cell lines generated in A, with H2B-TagBFP in blue, mNeonGreen-anillin in green and mRuby2-CAAX or mRuby2-actin in red. The scale bars are 25 μm. *This image was taken at a lower focal plane to show the actin filaments. C) Growth curves for the double- and triple-tagged HEK293 cell lines compared to unedited cells (WT) over 5 days, which is ∼5 population doubling times. D) A graph shows linescans comparing fluorescence intensity of mNeonGreen-anillin in single-tagged HEK293 cells (shown in black with standard deviation in grey) with double-tagged H2B-TagBFP; mNeonGreen-anillin HEK293 cells (shown in green).

### Generation of multi-tagged cell lines

It is highly desirable to visualize multiple proteins and/or cell components simultaneously. For example, co-expression of a tagged probe for chromatin can be used to measure the timing of cytokinesis. As a proof-of-concept, we generated HEK293 cell lines with multiple endogenous tags to visualize mNeonGreen-anillin with mRuby2 or TagBFP-histones, plasma membrane, actin and tubulin. We first tagged H2B (histone) with TagBFP and enriched a tagged cell population by FACS (Fig. 5A). Then we tagged anillin with mNeonGreen or the plasma membrane with mRuby2 in these enriched cell populations and isolated clonal cell lines (Fig. 5A, B). The double-tagged H2B-TagBFP; mNeonGreen-anillin cells were then used to generate triple-tagged cell lines by tagging the plasma membrane or β-actin with mRuby2 (mRuby2-CAAX and mRuby2-ACTB; Fig. 5B). We verified the overall health of these cell lines by measuring their growth rates and viability compared to non-edited HEK293 cells over five days (Fig. 5C). We found no major differences in the growth of multi-tagged cell lines compared to the non-edited cells, although H2B-TagBFP cells lagged behind for the first 2 days but recovered by day 3 (Fig. 5C). We also noticed that interphase H2B-TagBFP; mNeonGreen-anillin; mRuby2-actin cells displayed a different morphology than the non-edited cells or other cell lines, possibly because tagging actin could cause some disruption in actin dynamics. Despite this change, the growth rate of this cell line was similar to the others (Fig. 5C). Finally, we compared the intensity of mNeonGreen-anillin along the cortex of H2B-TagBFP; mNeonGreen-anillin cells compared to mNeonGreen-anillin cells. We found that the furrows in the double-tagged and single-tagged cells were well-aligned (Fig. 5D), showing that these tools can be used to measure protein levels in combination with other tagged cellular components, such as H2B-TagBFP.

## Discussion

In this study, we provide tools and protocols for the endogenous tagging of proteins and cell components to study cytokinesis. The majority of cytokinesis proteins have been studied using overexpressed transgenes in human cells, which may influence the interpretation of their function, and probes to visualize RhoA or its GEF Ect2 have been notoriously difficult to use. Endogenous tags can be useful not only to quantify endogenous proteins and measure changes in localization over time and/or in response to perturbation, but also can be useful to measure knockdown or when expressing mutant variants. We successfully tagged anillin, Ect2 and RhoA endogenously with mNeonGreen in HeLa and HEK293 cells, and then used the tagged HeLa cells to obtain valuable information regarding their localization by quantitative measurements. We found that their enrichment at the equatorial cortex occurred at similar times and correlated with when the ring would need to be fully assembled for ingression. We also found that anillin and RhoA were more broadly localized compared to Ect2. We then tagged anillin with mNeonGreen in additional cell lines: HCT116, HepG2 and MDCK cells. This permitted us to compare how cytokinesis occurs among very different cell types, based on differences in the timing, breadth and intensity of anillin localization. We found several key differences among the cell types that warrant further study. First, anillin was cortically localized in metaphase in HEK293, HCT116, and MDCK cells, but not in HeLa and HepG2 cells. It would be interesting to uncover why anillin is cortical in some cells and not others, which could reflect differences in the machineries required for cortical recruitment in the different cell types. We also observed that anillin was more broadly distributed along the equatorial cortex in HCT116 cells compared to the other cell types, while it was more narrowly distributed in HepG2 cells. Surprisingly, the accumulation of anillin in the equatorial peak did not correlate with ingression rates. For example, ingression was the fastest in HCT116 cells, and was the slowest in HepG2 cells, even though the enrichment of anillin was highest in HepG2 cells across all cell lines. We also observed two populations of HeLa cells in our cell line, where anillin accumulation was higher in one vs. the other, yet all cells showed similar breadth and ingression rates. Lastly, we successfully generated multi-tagged HEK293 cell lines to tag other components of the cell that would be useful for cytokinesis studies. These include using and expanding on the tools generated by the Allen Institute for Cell Science, and our constructs will also be made available on Addgene so that other researchers can use and improve on them. We endogenously tagged H2B with TagBFP with mNeonGreen-anillin and the plasma membrane (mRuby2-CAAX, integrated at the AAVS1 locus) or mRuby2-ACTB (β-actin). The general health of the different cell lines was comparable, suggesting that the tags did not interfere with function, however we did notice that interphase cells expressing mRuby2-actin had altered morphology. Although our efforts were focused on studying cytokinesis, several of these proteins may be of interest for studying other biological processes.

The generation of multi-tagged cell lines has been reported previously (Perez-Leal et al., 2021; Allencell.org). This is a promising strategy to visualize several proteins and markers without the need for transient transgene expression or fluorescent dyes. However, the culture time and clonal isolation required to introduce multiple tags can lead to cell senescence, and our method of using enriched populations of tagged cells for subsequent tagging vs. first isolating single clones drastically reduces the time needed for multi-tagging. Another concern is the compatibility of multiple tags for fluorescence microscopy. Some fluorescent signals have excitation/emission spectra that can overlap with filters in the adjacent channel, which can be a problem when combining two probes where one is weakly expressed requiring greater exposure times and the other is bright. Studies such as ours that provide additional fluorophore options with better separation in their excitation/emission spectra should overcome these issues. In addition, some tags fused to proteins such as β-actin or tubulin could potentially disrupt their assembly into polymers and/or dynamics. However, probes such as Lifeact or Sir-tubulin only report for polymerized actin and tubulin, respectively, and also could affect their dynamics. Thus, having more options to study monomers vs. polymers and/or that might have different impacts on assembly is highly desirable. Of the new endogenous tags generated in this work, anillin, Ect2 and RhoA, we found no unexpected cellular phenotypes or localization, suggesting that these tagged proteins were not disrupted significantly.

During this study, other groups reported the endogenous tagging of many proteins using self-complementing split fluorescent proteins (Mahdessian et al., 2021; opencell.czbiohub.org). This work validates our approach as they tagged Ect2 and RhoA on the same terminus and used the same sgRNA spacers as reported here. A caveat is that they did not look at their tagged proteins in the context of cytokinesis and it is known if they localize the same as the single fluorophores. The split-tagging approach can accelerate the generation of multiple endogenous tags in parallel. However, they are restricted to one cell type as it requires the generation of a parent cell line that expresses the core fragment of the split fluorescent protein to allow for tagging with the corresponding short fragment. Our tools (repair template and sgRNA) can be used to tag multiple proteins in any human cell line in one time.

With the cell lines generated during this work, we uncovered cytokinetic diversity across commonly used human and mammalian cell lines. Since most prior studies of human cytokinesis used HeLa cells, this work can lay the foundation for future studies of how mechanisms regulating cytokinesis vary with cell type. Overall, this study provides new tools to address new outstanding questions in the field of cytokinesis.

## Materials and Methods

### Cell culture

HEK293, HeLa and MDCK cells were cultured in Dulbecco’s modified Eagle medium (DMEM; Wisent) media supplemented with 10% Cosmic calf serum (CCS; Cytoviva). HCT116 cells were cultured in McCoy’s media (Wisent) supplemented with 10% CCS. HepG2 cells were cultured in Eagle’s minimum essential medium (EMEM; Wisent) supplemented with 10% Fetal bovine serum (FBS; Cytoviva). The cells were maintained in 10 cm dishes in incubators at 37°C with 5% CO_2_ as per standard protocols (Beaudet et al., 2017; Beaudet et al., 2020). For long-term storage, cells were resuspended in media (50% FBS, 40% DMEM or EMEM media and 10% DMSO) and stored in liquid nitrogen.

### Cloning

To generate the pX459-HypaCas9-mRuby2 CRISPR construct (Addgene #183872), we digested the pX459V2.0-HypaCas9 plasmid (Addgene #108294) with EcoRI (New England Biolabs) and used the GeneJET gel extraction kit (Thermo Fischer Scientific) to extract the backbone as per manufacturer’s protocols. To prevent re-ligation the backbone was dephosphorylated with Antarctic phosphatase (New England Biolabs) as per manufacturer’s instructions. The mRuby2-T2A-Puro insert was generated by amplifying the mRuby2 and Puromycin resistance gene with shared overlap and assembling them by SOEing (Splicing by Overlap Extension) PCR. This was done via first performing a Phusion PCR reaction (Thermo Fischer Scientific) without primers, then primers were added to the reaction for amplification by PCR as per manufacturer’s instructions. The mRuby2-T2A-Puro PCR product was then purified and digested with EcoRI for ligation into the dephosphorylated backbone. Clones were screened by PCR and validated by sequencing the insert region (Eurofins Operon).

The sgRNA spacer sequences were selected using the Benchling (Doench et al., 2016; Hsu et al., 2013) and CCTop (Stemmer et al., 2015) for the ANLN, ECT2, RHOA and MYH10 genes and the AAVS1 locus. Multiple spacers were selected for each target site. One of the AAVS1 spacer sequences was obtained from Oceguera-Yanez et al. (2016) and the spacer sequences for the HIST1H2BJ, ACTB, MYH10 and TUBA1B genes were obtained from the Allen Institute for Cell Science (Roberts et al., 2017; Allencell.org). All sgRNA spacer sequences tested are listed in Table S1. The sgRNA spacers were cloned into the pX459V2.0-HypaCas9 and the pX459V2.0-HypaCas9-mRuby2 plasmids using two complementary oligos containing the spacer sequence as follows (where (N)_20_ corresponds to the 20-nucleotide spacer sequence or its reverse complement): Forward oligo: 5’-CACCG(N)_20_-3’

Reverse oligo: 5’-AAAC(N)_20_-3’ Oligos were annealed then diluted in water, and ligated into the backbone (pX459V2.0-HypaCas9 or pX459V2.0-HypaCas9-mRuby2) which was pre-digested with BbsI (New England Biolabs) and gel purified. Two clones were selected for each sgRNA and sequenced to verify the spacer sequence.

To build the repair templates, the homology arms were first amplified from HEK293 genomic DNA extracted using the Qiagen DNeasy Blood and tissue kit, or plasmid DNA via a Phusion PCR reaction (Thermo Fischer Scientific). For genome sequences that were recalcitrant to PCR amplification, we used touchdown PCR conditions (Korbie & Mattick, 2008) with 1X GC buffer or 0.5X HF buffer instead of 1X HF buffer (Thermo Fischer Scientific). The fluorescent tags (mNeonGreen, mRuby2 and TagBFP) were amplified independently using primers that were designed to overlap with the homology arms (15-51 bp of homology). The primers used to clone the repair templates are listed in Table S2. The repair templates were assembled by SOEing PCR as described above, then blunt-end ligated into the pJET1.2 vector using the CloneJET PCR cloning kit (Thermo Fischer Scientific). Multiple clones were screened by PCR and sequencing.

For transgene expression, the coding sequence for anillin, Ect2 and Ect2(C-term) were cloned into the Golden Gate entry vector pYTK001 (Addgene #65108). They were fused to mNeonGreen (Allele Biotech) or mScarlet-I (Addgene #98839) and assembled into a custom Golden Gate expression vector (pGG). The Golden Gate assembly reaction was carried out using BsaI (New England Biolabs) following standard protocols (Lee et al., 2015). Isolated colonies were picked and screened by PCR and sequencing.

### Transfection, nucleofection and NHEJ inhibition

To improve editing efficiency, all cells were treated with NHEJ inhibitors NU7441 (Tocris Bioscience) and SCR7 (Xcess Biosciences) 4 hours before transfection and for 48 hours after transfection. HEK293, HeLa and HepG2 cells were seeded into 24-well plates to 60% confluency on the day of transfection. Transfections were carried out using Lipofectamine 3000 (Thermo Fischer Scientific) and P3000 (Thermo Fischer Scientific) reagent according to manufacturer’s instructions. Plasmids were introduced into HCT116 and MDCK cells using nucleofection with the 384-well HT Nucleofector and the SE cell line kit (Lonza) as per manufacturer’s instructions. The cells were nucleofected using the EN-113 program for HCT116 cells, and the CA-152 program for MDCK cells, and the amount of repair template and plasmid were adjusted per cell type.

For transient transgene expression, vectors [pGG-mNG-Anillin, pGG-mNG-Ect2, pGG-mScar-Ect2(C-term) and GFP-RhoA] were transfected into HeLa cells plated on coverslips to 60% confluency using Lipofectamine 3000 (Thermo Fisher Scientific) and P3000 reagent (Thermo Fisher Scientific) as per manufacturer’s instructions.

### Fluorescence-activated cell sorting

Tagged cells were isolated or enriched by FACS 7 to 14 days after transfection. The cells were resuspended thoroughly in FACS buffer comprised of 1 mM EDTA, 25 mM HEPES pH 7.0 and 1% FBS in PBS buffer, then passed through a 35 μm strainer to remove large cell clumps and transferred to 5 mL FACS tubes for sorting. Cells were sorted on a FACSMelody cell sorter (BD Biosciences) with gates were set to capture individual fluorescent cells. The fluorescent cells were then isolated into individual wells of a 96-well plate containing recovery media [media supplemented with 20% FBS and 1X Penicillin-Streptomycin (50 units/mL Penicillin and 50 μg/mL Streptomycin; Wisent)]. Alternatively, fluorescent cells were enriched by sorting 15,000 cells into a tube with recovery media. The enriched population was then resuspended in recovery media and seeded into wells in a 96-well plate. The cells were left to recover and media was supplemented as needed. For flow cytometry, cells were prepared using the same protocol and measured on different days after transfection. Flow cytometry data was acquired using the FACSMelody and analyzed with the R package CytoExploreR (Hammill, 2021).

### Genotyping

The first step of genotyping involved determining whether the tag was inserted into the locus using PCR. Clones were expanded by splitting cells from individual colonies into 48-well plates, and subsequently into 24-well plates. At this stage cells were harvested for PCR. The target locus was amplified using the Phire Plant Direct PCR mastermix (Thermo Fischer Scientific) in three junction PCR reactions as per manufacturer’s protocols. The primers used for these reactions are listed in Table S3. The three PCR reactions were designed to amplify the WT locus as well as the left and right junctions of the integration sites, and touchdown PCR was used to improve the quality of the product. 6-12 positive clones were then expanded into 10 cm dishes for sequencing and further use. When possible, homozygous clones carrying only tagged alleles were selected. Alternatively, heterozygous clones carrying a tagged allele and a WT allele were selected. Genomic DNA was extracted using the Qiagen DNeasy Blood and tissue kit, then used to amplify the target locus with Phusion polymerase (Thermo Fischer Scientific) as per manufacturer’s instructions by touchdown PCR. The genotype was verified by extracting PCR products and sequencing the tagged and non-tagged alleles.

### Microscopy

To collect still images of the endogenous tags and ectopic fluorescent protein expression, cells grown on 6-well plates were imaged 2 and 10 days after transfection on a Leica DMI6000B inverted epifluorescence microscope with filters for the appropriate wavelengths, a 20x/0.35 NA objective, an Orca R2 CCD camera (Hamamatsu) and Volocity software (PerkinElmer).

Single-cell-derived clones were initially screened for fluorescence and localization 7 to 10 days after FACS isolation using a Nikon Eclipse TS100 microscope equipped with a DS-Qi1Mc camera the 10x/0.25NA objective. Clones where every cell expressed a fluorescent signal at the expected cellular localization were selected for further screening.

To image cells during cytokinesis, tagged HeLa, HCT116, HepG2 and MDCK cell lines were seeded onto acid-etched round coverslips (25mm, No. 1.5) in 6-well plates and grown to 70% confluency. The coverslips were transferred to a magnetic chamber (Quorum) with 1 mL media before imaging. Tagged HEK293 cells were seeded directly onto 4-well μ-slides (Ibidi) for imaging. To visualize chromatin, Hoechst 33343 (Invitrogen) was added to the cells at a final concentration of 0.4 μM for 30 minutes prior to imaging. Imaging was performed using an inverted Nikon Eclipse Ti microscope (Nikon) equipped with a Livescan Sweptfield scanner (Nikon), Piezo Z stage (Prior), IXON 879 EMCCD camera (Andor), and 405, 488 and 561 nm lasers (100 mW, Agilent) using the 100x/1.45 NA objective. The cells were kept at 37°C and 5% CO_2_ during imaging in an INU-TiZ-F1 chamber (MadCityLabs). Images were acquired for 29 z-planes of 1 μm thickness every 1-2 minutes using NIS Elements software (Nikon, version 4.0).

To collect still images of the overexpressed transgenes, the coverslips were transferred to a magnetic chamber with 1 mL of media and imaged on a Leica DMI6000B inverted epifluorescence microscope with filters for the appropriate wavelengths, a 10x/0.25 NA objective, an Orca R2 CCD camera (Hamamatsu) and Volocity software (PerkinElmer), or a Nikon Eclipse TiE inverted epifluorescence microscope using LEDs in the appropriate wavelength with an Evolve 512 EMCCD camera (Photometrics) and NIS Elements acquisition software (Nikon) using the 60x/1.4 NA objective.

### Cell viability assay

To monitor growth of the different cell lines, we used the WST-8 cell proliferation assay kit (Cayman Chemical) as per manufacturer’s instructions. For each cell line, 10,000 cells were seeded into 96-well plates and cell viability was assayed on day 0, 1, 2, 3 and 5, corresponding to ∼4 population doubling times. For the assay, the electron mediator solution and the WST-8 developer reagent were mixed in equal parts, and added to each well and incubated at 37°C for 2 hours. After this, the absorbance at 450 nm was measured on a TECAN Infinite M200 plate reader. The assay was repeated in triplicate per cell line.

### Image analysis

Linescans were performed and measured for the different tagged cell lines using Fiji. Briefly, all images acquired using NIS Elements (Nikon) and Volocity (PerkinElmer) software were opened in Fiji (Version 2.3, NIH) and analyzed using a macro modified from Ozugergin et al. (2022). The macro first isolates the green channel from the movie file, subtracts background signal, and performs a bleach correction. The desired time point and Z slices are entered by the user, and the macro generates an average projection of the Z-stack. The user then manually traces a line along the cortex of the cell and a straight line intersecting the furrow. The macro then measures the fluorescence intensity of each pixel along the linescan, along with the length of the line and the position of the pixel intersecting the furrow. After analyzing cells with the macro, the data was imported into Excel (Microsoft) and Prism (Version 9.3, GraphPad) for further analysis.

### Statistical analysis

Box and whiskers plots were generated using Prism (Version 9.3, GraphPad) to show median value (central line), quartiles (box edges) and minimum and maximum values (whiskers). Statistical significance was tested using a Brown-Forsythe and Welch’s ANOVA, followed by multiple comparisons using Dunnett’s T3 test (Graphpad Prism version 9.3). Significance levels were defined as: p>0.5 non-significant (ns), * p≤0.05; ** p≤0.01; *** p≤0.001; **** p≤0.0001.

## Acknowledgements

We thank Andrew Cormier for his help in cloning the transgene expression vectors. We thank Dr. Chris Law and the Center for Microscopy and Cellular Imaging for guidance and help on image analysis. We thank Dr. Smita Amarnath, Nicholas D. Gold and the Center for Applied Synthetic Biology for training and use of the FACSMelody cell sorter. We also thank the McGill Flow Cytometry Innovation Platform for the sorting of BFP-tagged cells. We thank the Natural Sciences and Engineering Research Council of Canada (NSERC CREATE 511601-2018; NSERC Discovery Grant 04161-2017), the Fonds de Recherche Nature et Technologies (FRQNT B1X and B2X), Concordia University Research Chairs, and the National Research Council of Canada (NRCC) Disruptive Technology Solutions for Cell and Gene Therapy program for funding.

## Notes

### Competing Interest Statement

The authors have declared no competing interest.

