## Supplementary material for "Imaging tools generated by CRISPR/Cas9 tagging reveal cytokinetic diversity in mammalian cells"

**Table S1. List of sgRNAs used in this study.** The sequence of the sgRNAs tested for each target locus are listed, as well as their position relative to the site of integration. The oligos used to clone each sgRNA into the CRISPR plasmid (pX459V2.0-HypaCas9) are also shown. The preferred sgRNA for each site is highlighted in green.

| Gene/locus | sgRNA | Location | sgRNA sequence / PAM | Forward oligo | Reverse oligo | Orientation relative to gene | Distance relative to target site (bp) | Source |
|---|---|---|---|---|---|---|---|---|
| ANLN | 1 | N-terminus | TGGGGCGATGGATCCGTTTA/CGG | CACCGTGGGGCGATGGATCCGTTTA | AAACTAAACGGATCCATCGCCCCAC | Forward | +7 | This study |
| ANLN | 2 | N-terminus | GTCTCGTAGTCCGACGCCTG/GGG | CACCGGTCTCGTAGTCCGACGCCTG | AAACCAGGCGTCGGACTACGAGACC | Forward | -11 | This study |
| ANLN | 3 | N-terminus | GGACTACGAGACGATGGAAA/CGG | CACCGGGACTACGAGACGATGGAAA | AAACTTTCCATCGTCTCGTAGTCCC | Reverse | -33 | This study |
| ANLN | 4 | C-terminus | CATGGAAATTTCCCGGTTTA/AGG | CACCGCATGGAAATTTCCCGGTTTA | AAACTAAACCGGGAAATTTCCATGC | Reverse | +3 | This study |
| ANLN | 5 | C-terminus | TCAAAAACCTCTAGATAGCA/TGG | CACCGTCAAAAACCTCTAGATAGCA | AAACTGCTATCTAGAGGTTTTTGAC | Reverse | +21 | This study |
| ANLN | 6 | C-terminus | GATGCTTGCTACAAACCTAT/TGG | CACCGGATGCTTGCTACAAACCTAT | AAACATAGGTTTGTAGCAAGCATCC | Forward | -13 | This study |
| cfANLN | 1 | N-terminus | GGCGATGGACCCGTTTACCG/AGG | CACCGGGCGATGGACCCGTTTACCG | AAACCGGTAAACGGGTCCATCGCCC | Forward | +10 | This study |
| cfANLN | 2 | N-terminus | GGTAAACGGGTCCATCGCCC/CGG | CACCGGGTAAACGGGTCCATCGCCC | AAACGGGCGATGGACCCGTTTACCC | Reverse | +8 | This study |
| cfANLN | 3 | N-terminus | GCTTTCTCACCTCGGTAAAC/GGG | CACCGGCTTTCTCACCTCGGTAAAC | AAACGTTTACCGAGGTGAGAAAGCC | Reverse | -5 | This study |
| cfANLN | 4 | C-terminus | TATTGGGAAGCCTTAAACTG/TGG | CACCGTATTGGGAAGCCTTAAACTG | AAACCAGTTTAAGGCTTCCCAATAC | Forward | +4 | This study |
| cfANLN | 5 | C-terminus | CTTAAACTGTGGAAGTTTCC/AGG | CACCGCTTAAACTGTGGAAGTTTCC | AAACGGAAACTTCCACAGTTTAAGC | Forward | +15 | This study |
| cfANLN | 6 | C-terminus | GATGCTTGCTACAAACCTAT/TGG | CACCGGATGCTTGCTACAAACCTAT | AAACATAGGTTTGTAGCAAGCATCC | Forward | -13 | This study |
| cfANLN | 7 | C-terminus | ACAGTTTAAGGCTTCCCAAT/AGG | CACCGACAGTTTAAGGCTTCCCAAT | AAACATTGGGAAGCCTTAAACTGTC | Reverse | -9 | This study |
| ECT2 | 1 | N-terminus | AGTGTATTAACATCCACTAC/TGG | CACCGAGTGTATTAACATCCACTAC | AAACGTAGTGGATGTTAATACACTC | Forward | +26 | This study |
| ECT2 | 2 | N-terminus | CAAGCTAGTCCTCCCAGTAG/TGG | CACCGCAAGCTAGTCCTCCCAGTAG | AAACCTACTGGGAGGACTAGCTTGC | Reverse | +28 | This study |
| ECT2 | 3 | N-terminus | TATTAACATCCACTACTGGG/AGG | CACCGTATTAACATCCACTACTGGG | AAACCCCAGTAGTGGATGTTAATA | Forward | +30 | This study |
| ECT2 | 4 | C-terminus | GCCTTCCTTCCTTCTTTGAA/AGG | CACCGGCCTTCCTTCCTTCTTTGAA | AAACTTCAAAGAAGGAAGGAAGGCC | Forward | -45 | This study |
| ECT2 | 5 | C-terminus | CAGTTTCTTATTTGAGTATG/AGG | CACCGCAGTTTCTTATTTGAGTATG | AAACCATACTCAAATAAGAAACTGC | Reverse | +48 | This study |
| ECT2 | 6 | C-terminus | CTTCTCCTTTCAAAGAAGGA/AGG | CACCGCTTCTCCTTTCAAAGAAGGA | AAACTCCTTCTTTGAAAGGAGAAGC | Reverse | -51 | This study |
| ECT2 | 7 | C-terminus | ATTTCTATAATTTAAGATTT/TGG | CACCGATTTCTATAATTTAAGATTT | AAACAAATCTTAAATTATAGAAATC | Reverse | -64 | This study |
| ECT2 | 8 | C-terminus | AGAAGGAAGGAAGGCTGACA/AGG | CACCGAGAAGGAAGGAAGGCTGACA | AAACTGTCAGCCTTCCTTCCTTCTC | Reverse | +16 | This study |
| RHOA | 1 | N-terminus | AATCACCAGTTTCTTCCGGA/TGG | CACCGAATCACCAGTTTCTTCCGGA | AAACTCCGGAAGAAACTGGTGATTC | Reverse | +10 | This study |
| RHOA | 2 | N-terminus | GGCTGCCATCCGGAAGAAAC/TGG | CACCGGGCTGCCATCCGGAAGAAAC | AAACGTTTCTTCCGGATGGCAGCCC | Forward | +16 | This study |
| RHOA | 3 | N-terminus | TTTCAGCAATGGCTGCCATC/CGG | CACCGTTTCAGCAATGGCTGCCATC | AAACGATGGCAGCCATTGCTGAAAC | Forward | +6 | This study |
| H2BC11 | 1 | C-terminus | ACTCACTGTTTACTTAGCGC/TGG | CACCGACTCACTGTTTACTTAGCGC | AAACGCGCTAAGTAAACAGTGAGTC | Reverse | -5 | Allen Institute for Cell Science |
| ACTB | 1 | N-terminus | GCCGTTGTCGACGACGAGCG/CGG | CACCGGCCGTTGTCGACGACGAGCG | AAACCGCTCGTCGTCGACAACGGCC | Reverse | +19 | Allen Institute for Cell Science |
| MYH10 | 1 | N-terminus | GTTCTCTGCGCCATTGTAAA/TGG | CACCGGTTCTCTGCGCCATTGTAAA | AAACTTTACAATGGCGCAGAGAACC | Reverse | -6 | Allen Institute for Cell Science |
| MYH10 | 2 | N-terminus | TTTACAATGGCGCAGAGAAC/TGG | CACCGTTTACAATGGCGCAGAGAAC | AAACGTTCTCTGCGCCATTGTAAAC | Forward | +8 | Roberts et al., 2017 |
| MYH10 | 3 | N-terminus | TTGGATCGTTCCATTTACAA/TGG | CACCGTTGGATCGTTCCATTTACAA | AAACTTGTAAATGGAACGATCCAAC | Forward | -5 | This study |
| MYH10 | 4 | N-terminus | GGCGCAGAGAACTGGACTCG/AGG | CACCGGGCGCAGAGAACTGGACTCG | AAACCGAGTCCAGTTCTCTGCGCCC | Forward | +16 | Roberts et al., 2017 |
| TUBA1B | 1 | N-terminus | GATGCACTCACGCTGCGGGA/AGG | CACCGGATGCACTCACGCTGCGGGA | AAACTCCCGCAGCGTGAGTGCATCC | Reverse | -8 | Allen Institute for Cell Science |
| AAVS1 | 1 | - | GGGGCCACTAGGGACAGGAT/TGG | CACCGGGGGCCACTAGGGACAGGAT | AAACATCCTGTCCCTAGTGGCCCCC | Forward | +11 | Oceguera-Yanez et al., 2016 |
| AAVS1 | 2 | - | CTAGTGGCCCCACTGTGGGG/TGG | CACCGCTAGTGGCCCCACTGTGGGG | AAACCCCCACAGTGGGGCCACTAGC | Reverse | -12 | This study |
| AAVS1 | 3 | - | TGTCCCTAGTGGCCCCACTG/TGG | CACCGTGTCCCTAGTGGCCCCACTG | AAACCAGTGGGGCCACTAGGGACAC | Reverse | -7 | This study |

**Table S2. List of primers used for repair template cloning.** The primers used to amply the homology arms and the fluorescent markers to clone each repair template are shown. The different parts of the primer sequences are shown in different colors (annealing in green, tail in red, protein linker in dark blue and point mutations to abolish Cas9 cutting in light blue).

| **Construct** | **Terminus** | **Part** | **Orientation** | **Sequence** (annealing region / tail / protein linker / point mutations) |
|---|---|---|---|---|
| pODN-mNG-ANLN | N- | Left HA | Forward | ACAAAGCTACTGAAGGCCAACG |
| pODN-mNG-ANLN | N- | Left HA | Reverse | CGCCCCTGCCGTCGGACTACGAGACGATGCAATCGGTGGTTCAAATTCAGGAAGAGA |
| pODN-mNG-ANLN | N- | mNeonGreen | Forward | CCGACGGCAGGGGCGATGGTGAGCAAGGGCGAG |
| pODN-mNG-ANLN | N- | mNeonGreen | Reverse | ACTCACCTCCGTAAACGGATCCATGGAGCCTCCTGAACCTCCCTTGTACAGCTCGTCCATGCC |
| pODN-mNG-ANLN | N- | Right HA | Forward | ATGGATCCGTTTACGGAGGTGA |
| pODN-mNG-ANLN | N- | Right HA | Reverse | ACATCTCCCCGAAAGAATTACCAC |
| pODN-ANLN-mNG | C- | Left HA | Forward | AGCACAGTTTCCAGTACATGGAATG |
| pODN-ANLN-mNG | C- | Left HA | Reverse | AGGCTTTCCAATTGGCTTGTAGCAAGCATCAGGTTGCC |
| pODN-ANLN-mNG | C- | mNeonGreen | Forward | TTGCTACAAGCCAATTGGAAAGCCTGGAGGTTCAGGAGGCAGCATGGTGAGCAAGGGCGAG |
| pODN-ANLN-mNG | C- | mNeonGreen | Reverse | CCTCTAGATAGCCTTGAAATTTCCCAGCTTACTTGTACAGCTCGTCCATGC |
| pODN-ANLN-mNG | C- | Right HA | Forward | TAAGCTGGGAAATTTCAAGGCTATCTAGAGGTTTTTGATGTCATCTTAAGA |
| pODN-ANLN-mNG | C- | Right HA | Reverse | GGCGTTTAAAGGTGATAGGTGACTT |
| pODN-mNG-ECT2 | N- | Left HA | Forward | TCTAGCTTTTAAACTCTTCTTAAGTCCTGG |
| pODN-mNG-ECT2 | N- | Left HA | Reverse | CATGATTTGTATTCTTCTAAATCAGCTGAAAAAATAAA |
| pODN-mNG-ECT2 | N- | mNeonGreen | Forward | TTTTTATTCAAATTTTATTTTTTCAGCTGATTTAGAAGAATACAAATCATGGTGAGCAAGGGCGAG |
| pODN-mNG-ECT2 | N- | mNeonGreen | Reverse | TCGTAGAAGTTAGTACTGAATTCTCTGCCATGGAGCCTCCTGAACCTCCCTTGTACAGCTCGTCCATGCC |
| pODN-mNG-ECT2 | N- | Right HA | Forward | CCATGGCAGAGAATTCAGTACTAACTTCTACGACGGGAAGGACTAGCTTGGCAGACTCTT |
| pODN-mNG-ECT2 | N- | Right HA | Reverse | AGGACAACTTTCTTTTATCTTTATAACAGGGAC |
| pODN-ECT2-mNG | C- | Left HA | Forward | TAAATGCTTCATTTTACAACCTTGTTGATGAG |
| pODN-mNG-ECT2 | C- | Left HA | Reverse | AATCAAGTGAGTAGTAGACCGACTAAGCGTGTGACTCCGCCTCTCGAAAAAGCTAGGAAGGCTGACAAGGGAGG |
| pODN-mNG-ECT2 | C- | mNeonGreen | Forward | ACGCTTAGTCGGTCTACTACTCACTTGATTGGAGGTTCAGGAGGCAGCATGGTGAGCAAGGGCGAG |
| pODN-mNG-ECT2 | C- | mNeonGreen | Reverse | GACGTGTGTATATATTTGTATAGTTTGAGATATTAGTACCGATTCACTTGTACAGCTCGTCCATGCC |
| pODN-mNG-ECT2 | C- | Right HA | Forward | TGAATCGGTACTAATATCTCAAACTATACAAATATATACACACGTCTTACTCAAATAAGAAACTGACTTAAATGGTACTTG |
| pODN-mNG-ECT2 | C- | Right HA | Reverse | CTGACTATAAAAAGGCTATTGTGAAAGATATCACA |
| pODN-mNG-RHOA | N- | Left HA | Forward | TACCCTGCTTGACAAATTGTTTCCT |
| pODN-mNG-RHOA | N- | Left HA | Reverse | TGCTGAAACACAAAACACAGATATTACC |
| pODN-mNG-RHOA | N- | mNeonGreen | Forward | GCAGGTAATATCTGTGTTTTGTGTTTCAGCAATGGTGAGCAAGGGCGAG |
| pODN-mNG-RHOA | N- | mNeonGreen | Reverse | ACCAGTTTTTTGCGAATAGCTGCCATGGAGCCTCCTGAACCTCCCTTGTACAGCTCGTCCATGCC |
| pODN-mNG-RHOA | N- | Right HA | Forward | ATGGCAGCTATTCGCAAAAAACTGGTGATTGTTGGTGATGGA |
| pODN-mNG-RHOA | N- | Right HA | Reverse | GCCTGTAATCCCAGCTACTCTACT |
| pODN-mNG-cfANLN | N- | Left HA | Forward | TTTCAGCCCAACACGTCCAC |
| pODN-mNG-cfANLN | N- | Left HA | Reverse | CTCGACTGCGCGACGAC |
| pODN-mNG-cfANLN | N- | mNeonGreen | Forward | CGTCGTCGCGCAGTCGAGCCGCGGAGTCCGGGGCGATGGTGAGCAAGGGCGAG |
| pODN-mNG-cfANLN | N- | mNeonGreen | Reverse | GCTTTCTCACCTCGGTAAATGGATCCATGCTGCCTCCTGAACCTCCCTTGTACAGCTCGTCCATGCC |
| pODN-mNG-cfANLN | N- | Right HA | Forward | ATGGATCCATTTACCGAGGTGAGAAAGCCTG |
| pODN-mNG-cfANLN | N- | Right HA | Reverse | GTATGTTCTCAACAAGACTGGCCA |
| pODN-H2BC11-FP | C- | Left HA | Forward | CTTTGCCCAGCAGCTTGTTG |
| pODN-H2BC11-FP | C- | Left HA | Reverse | GGTGGCCACAGGAGGGT |
| pODN-H2BC11-FP | C- | Right HA | Forward | TAAACAGTGAGTTGGTTGCAAACTCT |
| pODN-H2BC11-FP | C- | Right HA | Reverse | CCTTTGCTTATATAGATATACCATAAAGGGAAAATAGT |
| pODN-H2BC11-mR2 | C- | mRuby2 | Forward | GGACCCTCCTGTGGCCACCATGGTGTCTAAGGGCGAAGAG |
| pODN-H2BC11-mR2 | C- | mRuby2 | Reverse | GATTGAGAGTTTGCAACCAACTCACTGTTTACTTGTACAGCTCGTCCATCCC |
| pODN-H2BC11-TagBFP | C- | TagBFP | Forward | GGACCCTCCTGTGGCCACCATGAGCGAACTGATCAAAGAGAACAT |
| pODN-H2BC11-TagBFP | C- | TagBFP | Reverse | GATTGAGAGTTTGCAACCAACTCACTGTTTAATTCAGTTTATGACCCAGCTTGCT |
| pODN-mR2-ACTB | N- | Left HA | Forward | GCTCGAGCGGCCGC |
| pODN-mR2-ACTB | N- | Left HA | Reverse | CATGGTGAGCTGCGAGAATAGCCGGGC |
| pODN-mR2-ACTB | N- | mRuby2 | Forward | GGCTATTCTCGCAGCTCACCATGGTGTCTAAGGGCGAAGAG |
| pODN-mR2-ACTB | N- | mRuby2 | Reverse | CATGGTACCGGAGCCGGCCTTGTACAGCTCGTCCATCCC |
| pODN-mR2-ACTB | N- | Right HA | Forward | GCCGGCTCCGGTACCATGGATGACGACATTGCAGCGCTCGTCGTCGACA |
| pODN-mR2-ACTB | N- | Right HA | Reverse | TACAGGGATAGCACAGCCTGG |
| pODN-mR2-MYH10 | N- | Left HA | Forward | AAGTGATGGAGCTTCCTCTGCT |
| pODN-mR2-MYH10 | N- | Left HA | Reverse | CATTGTAAATGGAACGATCCAAAAGCA |
| pODN-mR2-MYH10 | N- | mRuby2 | Forward | GCAATTGCTTTTGGATCGTTCCATTTACAATGGTGTCTAAGGGCGAAGAG |
| pODN-mR2-MYH10 | N- | mRuby2 | Reverse | CATAGGGATCCGCAGCTTCAGCTCCAGATCGCTGTACTTGTACAGCTCGTCCATCCC |
| pODN-mR2-MYH10 | N- | Right HA | Forward | AGCTGAAGCTGCGGATCCCTATGGCACAAAGGACAGGGCTGG |
| pODN-mR2-MYH10 | N- | Right HA | Reverse | TTAGTATCTGGCTTACATTTTTCCTCTCTACA |
| pODN-AAVS1-EGFP | - | Left HA | Forward | CTCAGTCTGGTCTATCTGCCTGG |
| pODN-AAVS1-EGFP | - | Left HA | Reverse | GGCCCCACTGTGTGGTCGAGGGGACAGATAAAAGTACCCAGA |
| pODN-AAVS1-EGFP | - | CMV-EGFP | Forward | CTCGACCACACAGTGGGGCCTAGTAATCAATTACGGGGTCATTAGTTCATAG |
| pODN-AAVS1-EGFP | - | CMV-EGFP | Reverse | GCTTTTCTGTCACTAAGCCTGTCCCTAGTAAAATTTAACGCGAATTTTAACAAAATATTAACGC |
| pODN-AAVS1-EGFP | - | Right HA | Forward | ACTAGGGACAGGCTTAGTGACAGAAAAGCCCCATCCT |
| pODN-AAVS1-EGFP | - | Right HA | Reverse | CCCCACAGTTGGAGGAGAATC |
| pODN-AAVS1-mNG | - | mNeonGreen | Forward | TGACTGACACCGGTCGCCACCATGGTGAGCAAGGGCGAG |
| pODN-AAVS1-mNG | - | mNeonGreen | Reverse | GCATGCATCGTACGTTACTTGTACAGCTCGTCCATGC |
| pODN-AAVS1-mNG-CAAX | - | mNeonGreen-CAAX | Forward | CGAATTCAAAGATGAGCAAAGATGGTAAAAAGAAGAAAAAGAAGTCAAAGACAAAGTGTGTAATTATGTAAAGCGGCCGCGACTC |
| pODN-AAVS1-mNG-CAAX | - | mNeonGreen-CAAX | Reverse | TCTTCTTTTTACCATCTTTGCTCATCTTTGAATTCGAAGCTTGAGCTCGAGATCTGAGTCCGGACTTGTACAGCTCGTCCATGCC |
| pODN-AAVS1-mR2 | - | mRuby2 | Forward | TGACTGACACCGGTCGCCACCATGGTGTCTAAGGGCGAAGAG |
| pODN-AAVS1-mR2 | - | mRuby2 | Reverse | GCATGCATCGTACGTTACTTGTACAGCTCGTCCATCCC |
| pODN-AAVS1-mR2-CAAX | - | mRuby2-CAAX | Forward | CGAATTCAAAGATGAGCAAAGATGGTAAAAAGAAGAAAAAGAAGTCAAAGACAAAGTGTGTAATTATGTAAAGCGGCCGCGACTC |
| pODN-AAVS1-mR2-CAAX | - | mRuby2-CAAX | Reverse | TCTTCTTTTTACCATCTTTGCTCATCTTTGAATTCGAAGCTTGAGCTCGAGATCTGAGTCCGGACTTGTACAGCTCGTCCATCCC |

**Table S3. List of primers used for genotyping.** The primers used to amplify each target locus for genotyping by PCR are shown.

| Gene/locus | Orientation | Sequence |
|---|---|---|
| ANLN | Forward | GCCGAGTCCGTCACTGG |
| ANLN | Reverse | TTCCTCCCTGAGTTTTATCTGTAGGAC |
| CfANLN | Forward | GGAGCAGTTTGGGAGCGTC |
| CfANLN | Reverse | ACACCCGCTGCGCAC |
| ECT2 | Forward | GAAATTGGTTGAGGGCAAAGGTG |
| ECT2 | Reverse | GCCTCCCTTTTCAAAGCTGC |
| RHOA | Forward | ATTCTTGTCTTGTTCCTGATCTTAGGG |
| RHOA | Reverse | ATTCTAACATGGAAAATGGCATCAGTTG |
| H2BC11 | Forward | TGGGCATCATGAATTCGTTTGTGA |
| H2BC11 | Reverse | TACTTCAGACCAAAAAACACAGTAGCA |
| ACTB | Forward | TAATAACGCGGCCGGCC |
| ACTB | Reverse | GAAGGAAAGGACAAGAAGCCCTG |
| MYH10 | Forward | TGAGGGCAAACCCATCAGAC |
| MYH10 | Reverse | TTTGATACTAGCTGCCTCAAAACCAT |
| TUBA1B | Forward | TGCAGTGAGCCTAGATGGCA |
| TUBA1B | Reverse | GAAGGTGTTGAAGGAGTCATCTCC |
| AAVS1 | Forward | GTCCACTTCAGGACAGCATGTT |
| AAVS1 | Reverse | CATCGTAAGCAAACCTTAGAGGTTCT |

### Figure S1

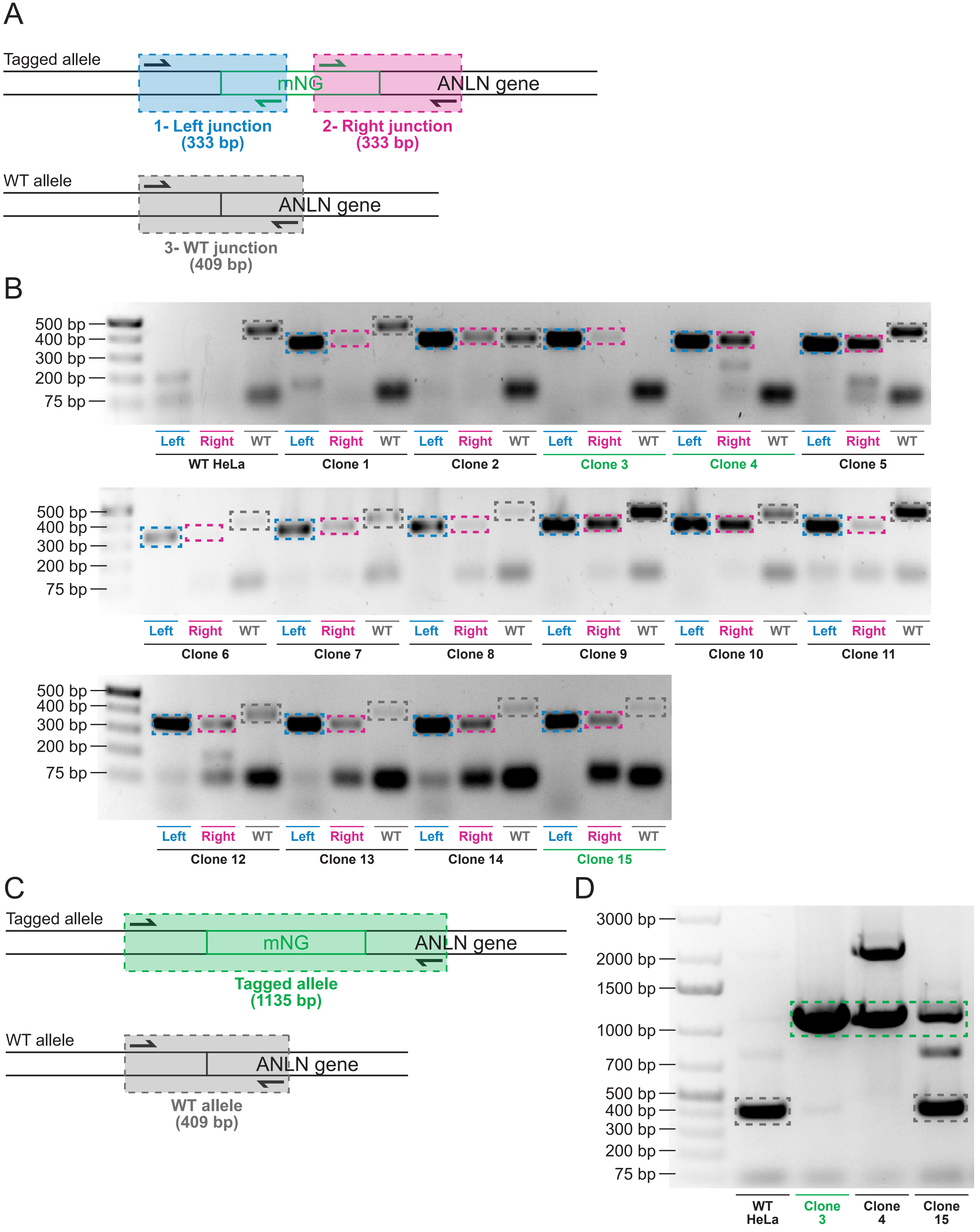

A
Tagged allele
mNG
ANLN gene
1- Left junction
(333 bp)
2- Right junction
(333 bp)
WT allele
ANLN gene
3- WT junction
(409 bp)
B
500 bp
400 bp
300 bp
200 bp
75 bp
Left Right WT
WT HeLa
Clone 1
Clone 2
Clone 3
Clone 4
Clone 5
Clone 6
Clone 7
Clone 8
Clone 9
Clone 10
Clone 11
Clone 12
Clone 13
Clone 14
Clone 15
C
Tagged allele
mNG
ANLN gene
Tagged allele
(1135 bp)
WT allele
ANLN gene
WT allele
(409 bp)
D
3000 bp
2000 bp
1500 bp
1000 bp
700 bp
500 bp
400 bp
300 bp
200 bp
75 bp
WT HeLa
Clone 3
Clone 4
Clone 15

**Figure S1. Screening and genotyping of mNeonGreen-anillin in HeLa cells.** A) The schematic shows how a junction PCR strategy was used to screen the mNeonGreen-anillin clones. Three PCR reactions (left junction, right junction and unedited, WT junction) were used to detect the presence of the tag and the non-tagged alleles. B) Gel images show junction PCR reactions for 15 mNeonGreen-anillin clones. The presence of each band is highlighted by colored boxes (blue for the left junction, pink for the right junction, and grey for the WT junction). The clones selected for further screening are indicated in green. C) A PCR genotyping strategy was used to validate the mNeonGreen-anillin HeLa cell line. The ANLN locus was amplified to detect the presence of tagged and non-tagged alleles. D) Gel images show genotyping of the mNeonGreen-anillin HeLa cell line. The presence of each band is highlighted by colored boxes (grey for non-tagged alleles and green for tagged alleles). The final clone selected for phenotyping (clone 3, highlighted in green) only carries tagged alleles.

### Figure S2

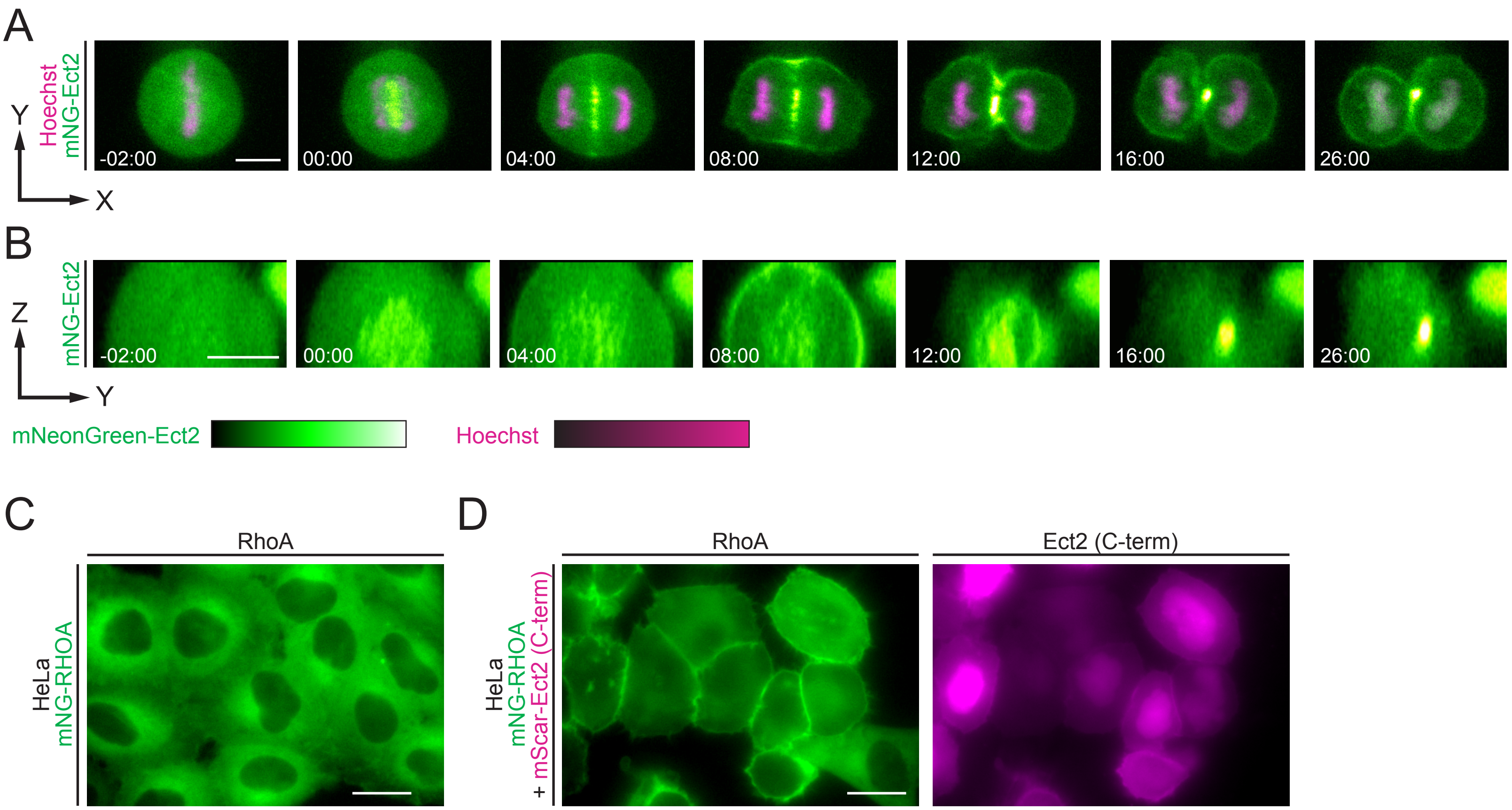


**Figure S2. Localization and regulation of mNeonGreen-Ect2 and mNeonGreen-RhoA in HeLa cells.** A) Timelapse images show endogenous mNeonGreen-Ect2 localization during cytokinesis. mNeonGreen is shown in green and DNA (stained by Hoechst) is in magenta. B) Timelapse images show an end-on ring view of mNeonGreen-Ect2 during cytokinesis. Times for A and B are shown relative to anaphase onset, and the scale bars are 10 µm. C) A representative image shows endogenous mNeonGreen-RhoA in interphase cells. D) Representative images show mNeonGreen-RhoA (left) in interphase cells co-expressing the C-terminus of Ect2 tagged with mScarlet-I (right). mNeonGreen is shown in green and mScarlet-I in magenta. The scales above show relative intensities. The scale bars are 20 µm.

### Figure S3

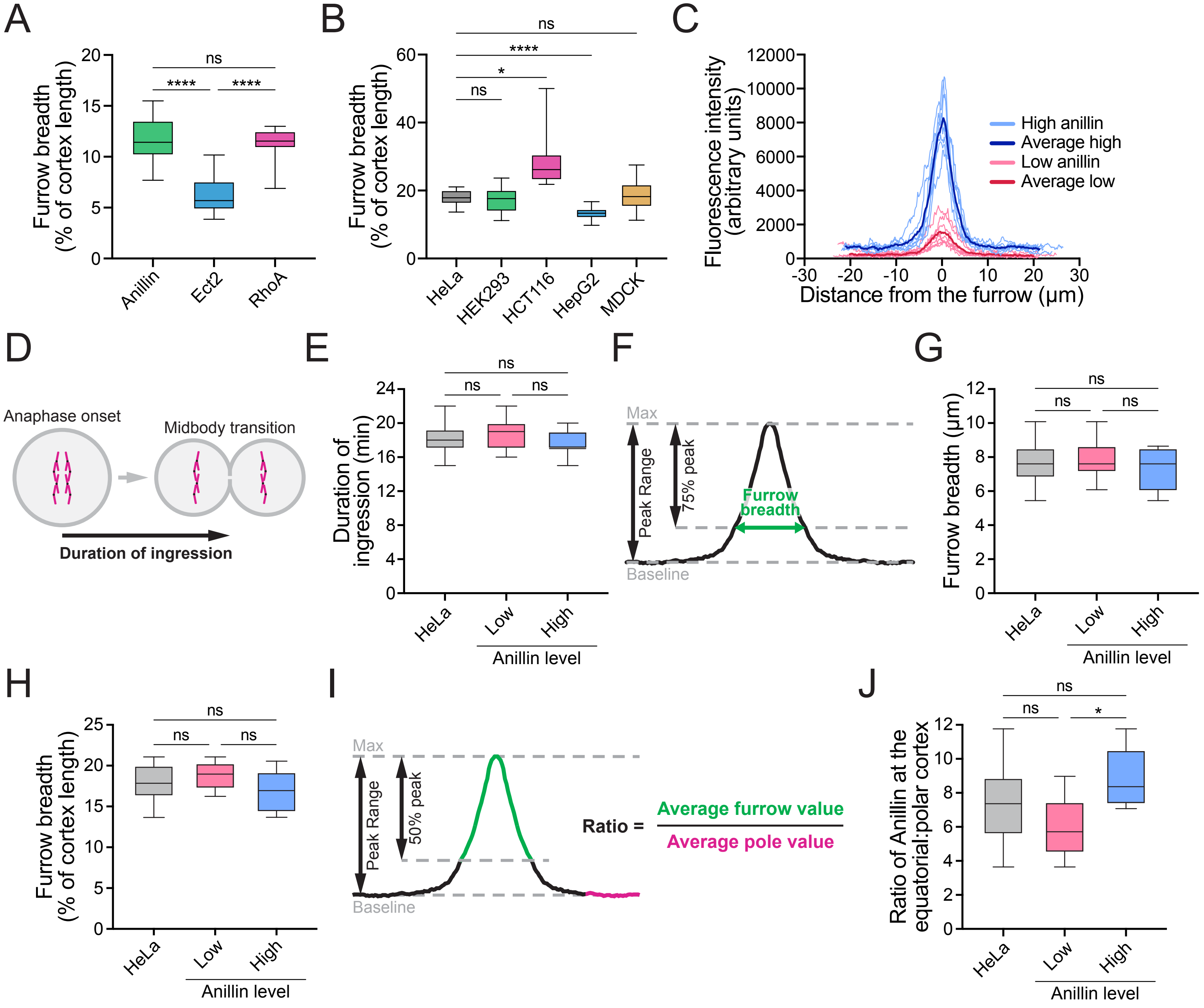
A
Furrow breadth (% of cortex length)
ns
****
****
Anillin
Ect2
RhoA
B
ns
****
*
ns
HeLa
HEK293
HCT116
HepG2
MDCK
C
Fluorescence intensity (arbitrary units)
High anillin
Average high
Low anillin
Average low
Distance from the furrow (μm)
D
Anaphase onset
Midbody transition
Duration of ingression
E
Duration of ingression (min)
ns
ns
ns
HeLa
Low
High
Anillin level
F
Max
Peak Range
75% peak
Furrow breadth
Baseline
G
Furrow breadth (μm)
ns
ns
ns
HeLa
Low
High
Anillin level
H
Furrow breadth (% of cortex length)
ns
ns
ns
HeLa
Low
High
Anillin level
I
Max
Peak Range
50% peak
Baseline
Ratio =
Average furrow value
Average pole value
J
Ratio of Anillin at the equatorial:polar cortex
ns
ns
*
HeLa
Low
High
Anillin level

**Figure S3. Characterization of endogenous mNeonGreen-anillin, Ect2 and RhoA localization during cytokinesis.** A) Box plots show the breadth of mNeonGreen-anillin, -Ect2 and -RhoA in HeLa cells relative to cortical length. B) Box plots show the breadth of mNeonGreen-anillin in HeLa, HEK293, HCT116, HepG2 and MDCK cells relative to cortical length. C) A graph shows two sets of linescans to highlight the differences in fluorescence intensity of mNeonGreen-anillin along the cortex of 'high' (blue, n=8) and 'low' (red, n=8) expressing HeLa cells. Individual cells are shown in light colors and the average for each population is shown in dark colors. D) A schematic shows how ingression time was measured. E) A boxplot shows the duration of ingression in high- and low- expressing mNeonGreen-anillin HeLa cells compared to combined populations. F) A schematic shows how the breadth at the equatorial cortex was calculated for G and H. G) A boxplot shows the breadth of mNeonGreen-anillin in high- and low- expressing HeLa cells compared to combined populations. H) A boxplot shows the breadth of mNeonGreen-anillin relative to cortical length in high- and low- expressing HeLa cells compared to combined populations. I) A schematic shows how the ratio of protein at the equatorial cortex relative to the polar cortex was calculated for J. J) A boxplot shows the ratio of mNeonGreen-anillin at the equatorial cortex vs. polar cortex in high- and low- expressing HeLa cells compared to combined populations. Bars for all graphs show min and max values, and statistical significance was determined by one-way ANOVA (ns, not significant; * $p \leq 0.05$; ** $p \leq 0.01$; *** $p \leq 0.001$; **** $p \leq 0.0001$). Box plots in A, B, E, G, H and J show the median line, quartile box edges and minimum and maximum whiskers.

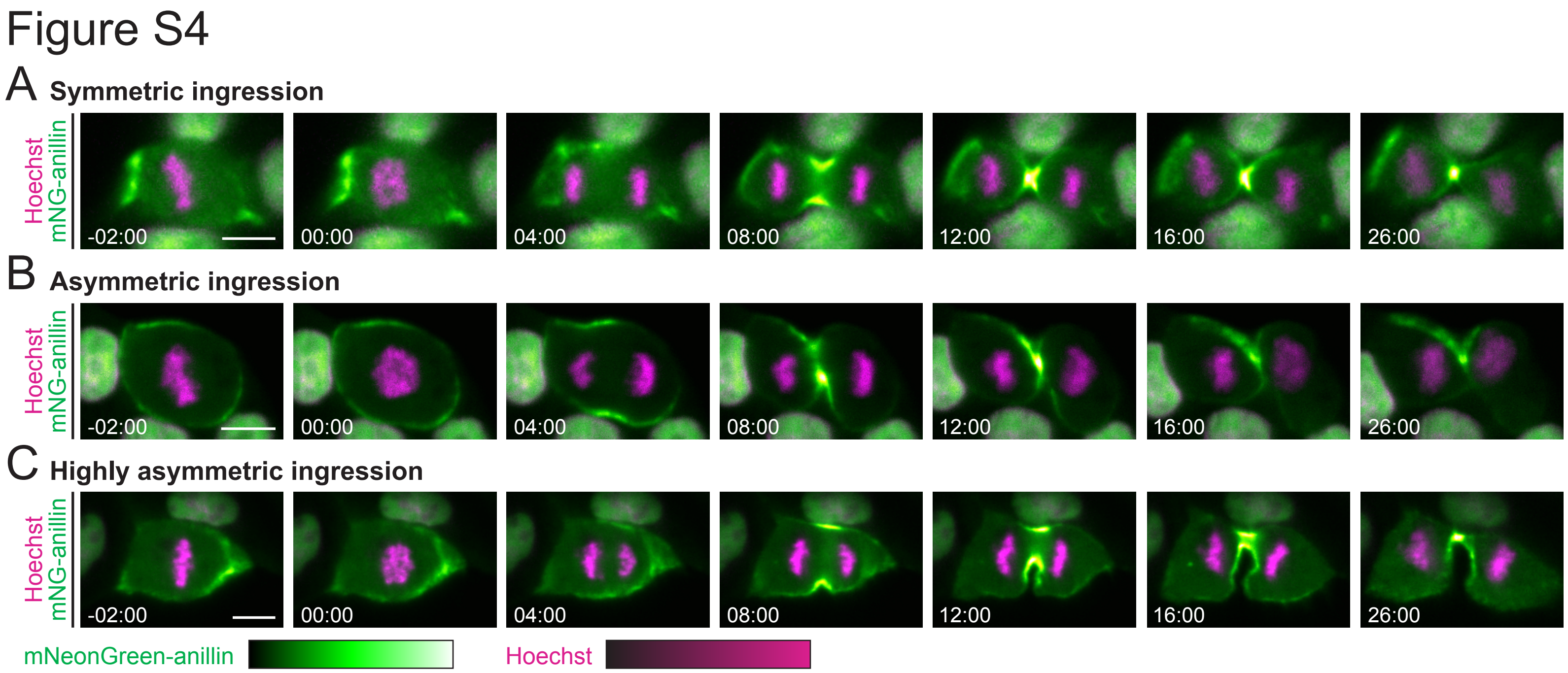


**Figures S4. Ingression occurs with variable symmetry in MDCK cells.** Timelapse images show endogenous mNeonGreen-anillin localization in different MDCK cells during cytokinesis where ingression appeared to be more symmetric (A), asymmetric (B), or highly asymmetric (C). mNeonGreen is shown in green, and DNA (stained by Hoechst) is in magenta, and relative intensity is shown in the corresponding scales. Times are from anaphase onset, and the scale bars are 10 μm.
